# Topologically controlled circuits of human iPSC-derived neurons for electrophysiology recordings

**DOI:** 10.1101/2021.12.10.472063

**Authors:** Sophie Girardin, Blandine Clément, Stephan J. Ihle, Sean Weaver, Jana B. Petr, José C. Mateus, Jens Duru, Csaba Forró, Tobias Ruff, Isabelle Fruh, Matthias Müller, János Vörös

## Abstract

Bottom-up neuroscience, which consists of building and studying controlled networks of neurons *in vitro*, is a promising method to investigate information processing at the neuronal level. However, *in vitro* studies tend to use cells of animal origin rather than human neurons, leading to conclusions that might not be generalizable to humans and limiting the possibilities for relevant studies on neurological disorders. Here we present a method to build arrays of topologically controlled circuits of human induced pluripotent stem cell (iPSC)-derived neurons. The circuits consist of 4 to 50 neurons with mostly unidirectional connections, confined by microfabricated polydimethylsiloxane (PDMS) membranes. Such circuits were characterized using optical imaging and microelectrode arrays (MEAs). Electrophysiology recordings were performed on circuits of human iPSC-derived neurons for at least 4.5 months. We believe that the capacity to build small and controlled circuits of human iPSC-derived neurons holds great promise to better understand the fundamental principles of information processing and storing in the brain.

## 1 Introduction

A major unanswered question in neuroscience is how the human brain processes and stores information. Unraveling the basic principles of neural computation would not only advance the fundamental understanding of the brain but could also help to elucidate the mechanisms behind and treatment of neurological diseases. Further, such findings can provide guidance for studies of neural regeneration and improve brain-machine interfaces for neuroprosthetics ^1^. It has been established that the primary information processing cells in mammals are neurons, which transmit information through electrical signals ^2^. However, electrical signalling at the neuronal level is difficult to investigate *in vivo* due to the complex and densely packed architecture of the brain and the limited resolution of the experimental tools available ^3^. An alternative and promising approach to gain knowledge about neural information processing is “bottom-up” neuroscience, which consists of engineering and studying elementary *in vitro* networks of neurons to understand gradually more complex systems ^1,4,5^. This approach could provide the technological tools needed to analyze how the structure and geometry of a controlled assembly of neurons affect its functional electrical activity.

*In vitro* networks of neurons can be engineered through two main approaches: surface patterning and physical confinement of the neurons. Surface patterning consists of depositing specific molecules on a substrate to define cell-attractive and cell-repellent areas. Patterning is commonly achieved using techniques such as microcontact printing ^6–9^ or photolithography ^10–12^. However, neurons that connect together exert forces on each other leading to clustering and gradual changes in the network architecture ^13^. In addition, coatings are degraded by the cells over time making it challenging to keep consistently patterned cultures over the long term. Since neurons typically take a week or more to become electrically active and functionally mature *in vitro* ^14,15^, it is desirable to build networks that are stable over several weeks to be able to investigate their functional electrical activity. Therefore, an alternative and more adopted method to engineer biological neuronal networks is the use of three-dimensional microfabricated structures to spatially confine cell bodies ^16^. The most widely used material to build such microstructures is polydimethylsiloxane (PDMS), which is biocompatible, transparent, and easy to process. An additional advantage of microstructures compared to surface patterning techniques is that they can be designed to directionally guide axons between groups of neurons ^17–20^. As axons transmit action potentials from one neuron to the next, controlling the direction of growth of axons should allow defining the main direction of information flow in a network. Action potential transmission in microstructures can be measured by aligning and bonding the PDMS structures to microelectrode arrays (MEAs) ^21–23^.

An important, yet seldom discussed, consideration for bottomup neuroscience is the source of cells chosen to build *in vitro* neuronal networks. There are three possible sources of neurons: immortalized neuronal cell lines, primary neurons, and stem cell-derived neurons. Each can originate from either model animals (mostly rodent) or humans. The first source of cells, immortalized neuronal cell lines, is derived from tumours. These cells are easy to culture and to expand, but are ill-suited for building *in vitro* neuronal networks because they usually present altered physiology and abundant genetic aberrations ^24^. The second cell source, primary cells, presents more physiologically relevant characteristics ^25^. Rodent primary neurons, especially from rats, have been extensively used in bottom-up neuroscience investigations ^26–28^. However, as rat primary neurons are dissociated from brain cells of embryos or pups, they result in a heterogeneous cell population ^29^ and might lead to variations across experiments. In addition, new animals must be sacrificed for each culture, which is incompatible with concerted efforts to reduce the number of animals used in scientific experiments, in particular the “3R initiative” ^30^. Finally, due to inter-species differences the conclusions made with rodent primary neurons might not be generalizable to humans, especially when investigating neurological disorders ^31–33^. This is well illustrated by the fact that in dissociated cultures, maturation time and network activity differ significantly between cultures of rat and of human primary neurons ^34^. Investigations using adult brain slices from several mammalian species have revealed that human cortical pyramidal neurons have a unique biophysical composition compared to all other species, presenting a different dendritic physiology and lower conductances than expected for their size ^35^. Considering all these elements, neurons of human origin should be used for *in vitro* investigations to generate more conclusive data. However, access to adult human primary brain tissue that can be dissociated for *in vitro* cell culture is limited ^36^ and access to embryonic human brain tissue raises ethical questions, since such tissues originate from aborted human fetuses ^37^.

The third possible cell source consists in using human stem cell-derived cells to generate differentiated cell types. The two main types of stem cells are embryonic stem cells (ESCs) and induced pluripotent stem cells (iPSCs), both of which are self-renewing and pluripotent. On the one hand, access to human ESCs is restricted and legally regulated because they mostly originate from discarded *in vitro* fertilized human embryos ^38^. On the other hand, iPSCs are widely available because they are obtained by reprogramming adult somatic cells through the addition of small molecules or the forced expression of genes coding for specific transcription factors ^39^. Adult somatic cells are easy to obtain, for example through a skin biopsy or blood sample ^40^. Several methods now exist to reprogram iPSCs into neurons, many of which allow differentiated neurons to be cryopreserved. This presents two major advantages: first, neurons coming from the same source can be used across numerous experiments, which should decrease the inter-experiment variance; second, laboratories that do not have the required biological facilities and expertise to produce iPSCs themselves now have access to iPSC-derived neurons, either commercially or through collaborations. Overall, iPSC-derived neurons have the potential to lead to more human-relevant conclusions, to provide homogeneous and tailorable differentiated cell types, to reduce the use of animals in experiments and to be easily accessible across laboratories. For all of these reasons, we consider iPSC-derived neurons to be the most suitable cell source for many bottom-up neuroscience investigations.

A reliable method to differentiate human iPSC into neurons is through the overexpression of the gene Neurogenin 2 (Ngn2), as first reported by Zhang *et al*. ^41^ and later refined by several groups ^15,42–44^. Compared to previous methods, this method is fast, has a high conversion efficiency, and produces neurons with properties independent of the starting iPSC line ^44^. Neurons obtained through the overexpression of Ngn2 are termed “induced” neurons, or iNeurons, and present properties similar to those of cortical glutamatergic excitatory neurons ^41^. Such iNeurons have recently been used in microfluidic multi-compartment chambers (“Taylor” chambers) for drug screening applications ^45^ and together with dopaminergic and inhibitory neurons to study neuronal subtype connections ^46^. iPSC-derived cortical neurons obtained through other methods than Ngn2 overexpression have also been used in combination with Taylor chambers to study axonal damage ^47^, *α*-synuclein propagation ^48^, long-term development ^49^, and connections between neurons of the peripheral and central nervous systems ^50^. However, in all of these studies, the number of neurons inspected was on the order of 10^4^ to 10^5^ neurons per compartment. We believe that to reduce the variability that arise from the complexity of such networks and to get a more reproducible network behavior, it is necessary to be able to build networks with a lower number of neurons, in the range of single to tens of cells per compartment. Neuronal cultures at such low density are challenging to maintain and require protocol optimization.

Here we report the use of human iNeurons to build biological neuronal circuits, each composed of less than 50 cells and with predominantly unidirectional connections. The topology of the neuronal circuits is controlled using thin microfabricated PDMS membranes, which can be placed on top of MEAs to record electrophysiology data from the circuits. The engineered neuronal circuits were also characterized using fluorescent stains. We optimized the culture protocol to obtain reasonable cell survival despite the low seeding density and recorded spontaneous electrical activity of some of the circuits for up to 133 days *in vitro* (DIV). All in all, the technology presented here provides a modular platform to build and deconstruct circuits of human neurons over several months, with the potential to investigate the fundamental biophysics of information processing, plasticity mechanisms, and neurophysiological disorders.

## 2 Materials and Methods

### 2.1 PDMS microstructures

Polydimethylsiloxane (PDMS) microstructures were designed in Python using the GDScad package, based on a template from Forró *et al*. ^20^ shown on Fig. 1a. A typical microstructure contained a set of 15 circuits (see Fig. 1a). They were fabricated on a 4-inch wafer by Wunderlichips (Switzerland) using a standard soft lithography process ^20^. The resulting PDMS membrane is a two-layer structure with a first layer with a height of about 200 μm, with cylindrical nodes with a diameter of either 100 or 170 μm; and a second layer with a height of about 4 μm, connecting the holes through narrow microchannels (Fig. 1b). Before use, the microstructures were cut out of the PDMS wafer with a scalpel, cleaned of any dust using Scotch tape and left on a clean glass slide until use. In rare cases, due to the microfabrication process, a thin layer of PDMS remained on top of one or more of the nodes of a microstructure, later preventing iNeurons from falling inside. Such nodes were identified during image post-processing and excluded from the analysis.

**Fig. 1.**
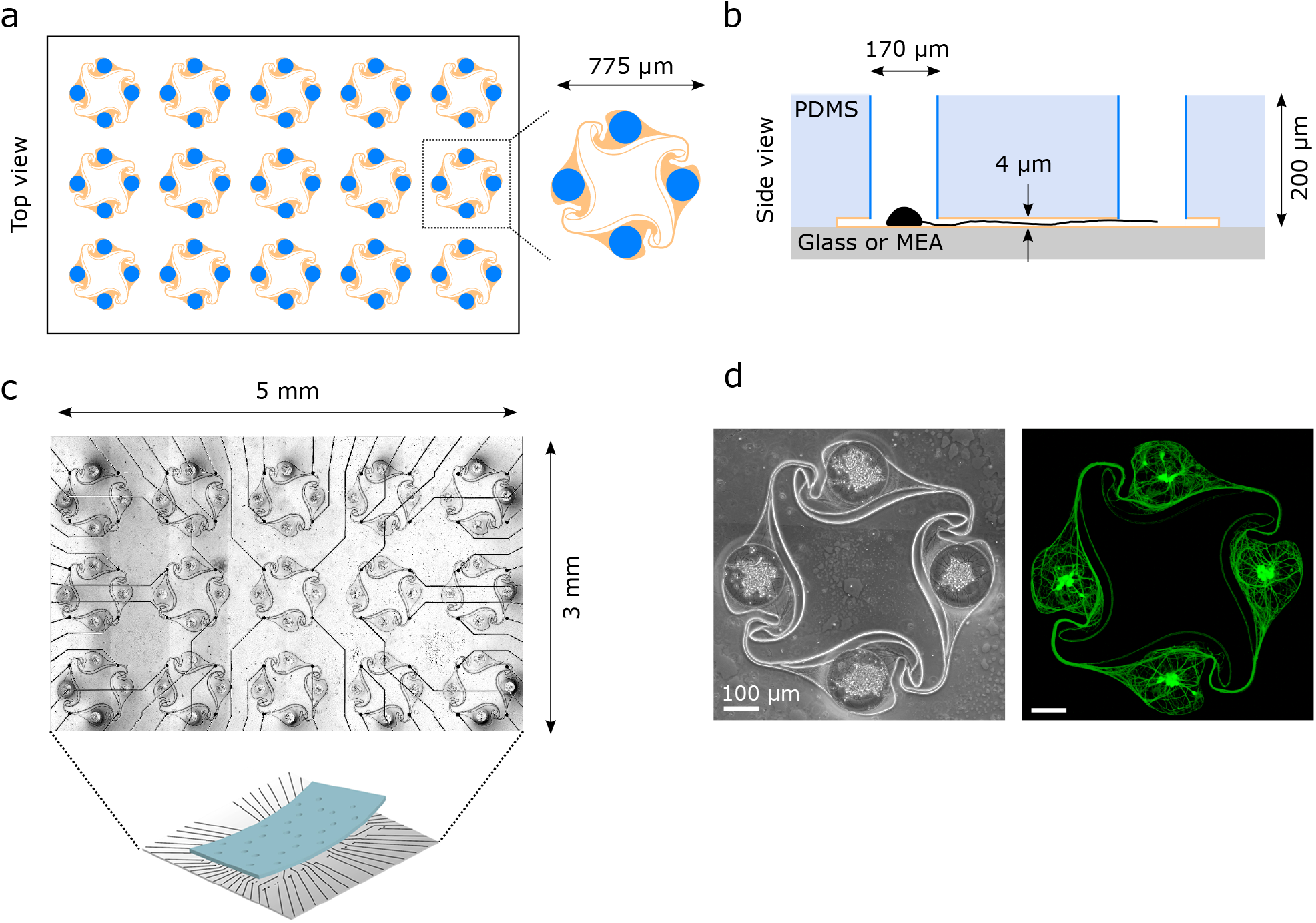
Overview of the PDMS microstructures used to build circuits of iNeurons with controlled axon guidance. (a) Top view of the layout of a typical PDMS microstructure, consisting of 15 circuits, with a zoom-in on one of the circuits. A circuit consists of four nodes (blue) connected by narrow microchannels (orange). The “stomach” shape of the channels allows for axon guidance, resulting in mostly unidirectional, clockwise connections between the nodes (see Fig. 6). (b) Schematic side view of two nodes (blue) connected by a microchannel (orange) where an axon is growing [not to scale]. The microchannels are too low for the soma to migrate into, resulting in the physical confinement of the soma in the nodes. (c) Micrograph of a PDMS microstructure with 15 circuits aligned to the 60 electrodes of a MEA. One electrode is positioned under each of the four narrow microchannels of a 4-node circuit, allowing to record from the axon bundle passing on top. (d) Example of a circuit of iNeurons cultured in a PDMS microstructure: phase-contrast (left) and fluorescently labelled iNeurons (right, stained with Calcein AM). The soma can be identified as the brighter spots visible in the center of each node.

### 2.2 Substrate preparation

All imaging experiments were performed using glass bottom 35-mm diameter dishes (HBST-3522T, WillCo Wells) as a substrate, unless mentioned otherwise. A typical phase contrast and a fluorescent image of a formed iNeuron network is shown in Fig. 1d. All electrophysiological experiments were performed on 60-electrode microelectrode arrays (60MEA500/30iR-Ti-gr, Multi Channel Systems).

#### 2.2.1 Glass bottom dish preparation

30-mm diameter coverslips (Menzel glass, selected #1.5, ThermoFisher) were cleaned with acetone, isopropanol, and ultrapure water (Milli-Q, Merck-MilliPore) before being blow dried with nitrogen. The glass bottom dishes were then mounted according to the manufacturer’s instructions.

The dishes were plasma cleaned for 2 min (18 W PDC-32G, Harrick Plasma) and coated with 300 μL per dish of 0.1 mg/mL PDL (P6407, Sigma Aldrich) in PBS (10010-023, ThermoFisher). After 45 min, dishes were rinsed three times with PBS and left in ultrapure water. The water left in the glass bottom dish was then aspirated and two PDMS microstructures were placed in the dish using tweezers. After inspecting the dishes under a stereo microscope to check if the structures were lying flat against the bottom of the dish, they were blow dried and placed in a desiccator for 10 min to ensure proper adhesion of the PDMS membrane to the glass. 2 mL of warm PBS was added to the dish before placing it in the desiccator for at least one hour to remove the air trapped in the narrow channels of the microstructures. Dishes were then stored in PBS at 4 °C for up to three days before cell seeding.

Laminin (11243217001, Sigma Aldrich) was optionally used as a secondary coating by adding it to a dish that already contained the PDMS microstructure at a concentration of 10 μg/mL in PBS at 37 °C for 2 h, before rinsing it once with culture medium and seeding cells on the sample.

#### 2.2.2 MEA preparation

Microelectrode arrays (MEAs) can be reused across several experiments. When reusing a MEA, it was first immersed in a solution of 4 % Tergazyme (Alconox, 1304-1) for 24 h to remove cell culture and proteins, then kept in ultrapure water until reuse. On the day of substrate preparation, it was cleaned three times with 0.2 % w/V sodium dodecyl sulfate (SDS, L3771, Sigma Aldrich), ultrapure water, ethanol and ultrapure water again, before being blow dried with nitrogen. No cleaning steps were performed for new MEAs.

MEAs were oxygen plasma cleaned for 2 min and coated with 250 μL of 0.1 mg/mL PDL in PBS for 45 min. This was followed by three subsequent rinses with PBS, before leaving the MEAs in ultrapure water. The water left in the MEA was aspirated away, leaving a thin layer of liquid, and a microstructure was placed in the dish using tweezers. The tweezers were used to carefully align the microchannels to the electrodes of the MEA (see Fig. 1c). The MEA was then blow dried and placed in a desiccator for 10 min to ensure proper adhesion of the PDMS to the glass. 2 mL of warm PBS was then added to the MEA and it was placed in the desiccator for at least one hour to remove air trapped in the channels of the microstructures. MEAs were then stored in PBS at 4 °C overnight before cell seeding.

#### 2.2.3 Glass bottom well plate

For open cultures of iNeurons, a glass bottom 48-well plate (P48G-1.5-6-F, Mattek) was used as a substrate. It was plasma cleaned for 2 min and coated with 0.1 mg/mL PDL in PBS for 45 min before being rinsed 3 times with PBS and left in ultrapure water.

### 2.3 iNeuron culture

#### 2.3.1 iPSC differentiation

Human iPSCs were generated following a previously published protocol ^51^ and transfected with a doxycycline-inducible Neurogenin-2 (Ngn2) gene. Differentiation into neurons was induced by a 3-day exposure to doxycycline as reported in Russell *et al*. ^52^. Differentiated iNeurons were then cryogenized as aliquots of 1 · 10^6^ to 8 · 10^6^ cells in heat inactivated FBS containing 5 % DMSO. Cryogenized aliquots of iNeurons were kindly provided by Novartis and stored in liquid nitrogen until use.

#### 2.3.2 NBD medium

The culture medium used with iNeurons was Neurobasal differentiation medium (NBD). NBD was prepared freshly by adding 1 mL of B27 supplement (17504-044, ThermoFisher), 0.5 mL of N2 supplement(17502-048, ThermoFisher), 50 μL of brain-derived neurotrophic factor (BDNF, 10 μg/mL, 450-10, PeproTech) and 50 μL of glial-derived neutrophic factor (GDNF, 10 μg/mL, 450-02, PeproTech) to 50 mL of Neurobasal medium (NeuroBasal medium (21203-049) with an added 1 % GlutaMAX (35050-061) and 1 % Pen-Strep (15070-063, all from ThermoFisher).

#### 2.3.3 iNeuron seeding and culture

About 2 h prior to cell seeding, the PBS contained in the substrates (glass bottom dish, MEA or well plate) was replaced with 1 mL of NBD. The substrates were placed in an incubator (37 °C, 5 % CO_2_, Steri-Cycle 371 CO_2_ Incubator, Thermo Fisher Scientific) until seeding.

An iNeuron aliquot was taken out of the liquid nitrogen and put at 37 °C to thaw rapidly. The 1 mL thawed cell solution was transferred dropwise into 4 mL of warm NBD and centrifuged for 5 min at 1000 rpm. The supernatant was aspirated and cells were resuspended at a concentration of 1 · 10^6^ cells/mL. The cell solution was passed through a 40 μm strainer (CSS013040, BioFilJet) and counted using a cell counter (Cell Countess, Invitrogen).

A volume containing the target cell number was pipetted onto the substrate (30 to 65 k cells/cm^2^ for PDMS microstructures substrates; 300 k cells/cm^2^ on the glass bottom well plates). After 10 min, the solution was mixed by pipetting to increase the number of iNeurons in the PDMS nodes. A complete medium exchange was done 1 h after seeding to remove dead cells. For the laminin-supplemented experiments, laminin was added to the medium at this stage, to a final concentration of 1 to 10 μg/mL. A half medium change was performed two to three times a week, with optional addition of laminin to the medium during the first week of medium change. In all experiments, the day of iNeuron thawing and seeding was considered as DIV 0.

### 2.4 Staining and imaging

#### 2.4.1 CMFDA staining

CMFDA (1 mM in DMSO, CellTracker Green CMFDA Dye, C7025, ThermoFisher) and ethidium homodimer-1 (2 mM in DMSO, L3224, ThermoFisher) were added directly to the cell medium to a final concentration of 1 μM each. The sample was incubated for 30 min before replacing the medium with fresh, warm NBD.

#### 2.4.2 Live-dead and Hoechst staining

A solution of 2 μM of Calcein-AM and 8 μM of ethidium homodimer-1 (both from L3224, ThermoFisher) in DPBS (14190-144, ThermoFisher) was incubated with the sample for 12 min. The same volume of a solution of 2 μM of Hoechst 33342 (H3570, ThermoFisher) was added to the sample and incubated for another 8 min. The sample was then carefully washed once with DPBS and left in warm DPBS for imaging.

#### 2.4.3 Image acquisition

A confocal laser scanning microscope (FluoView 3000, Olympus) was used to image the stained cultures. Three to four channels were typically acquired: 405 nm (Hoechst), 488 nm (Calcein AM or CMFDA), 561 nm (ethidium homodimer-1) and phase contrast brightfield images.

#### 2.4.4 Image analysis

Microscope images were processed using Fiji ^53^. Importantly, due to their size, stained soma are brighter and thus more visible than axons on microscopy images. To enhance the intensity of the axons compared to the soma, a pixel logarithm operator was applied to all the representative fluorescent images shown in the figures of this paper. The brightness and contrast were manually adjusted to suppress background fluorescence.

#### 2.4.5 Statistical tests

Boxplots were used to represent the data. The interquartile range (IQR) was calculated as the difference between the 3rd quartile (Q3) and the 1st quartile (Q1). On the boxplot: the bottom whisker is the closest data above Q1 - (1.5 x IQR); the coloured part of the box is bounded by Q1 and Q3; the middle horizontal black bar indicates the median; the top whisker is the closest data below Q3 + (1.5 x IQR). Outliers are indicated as single points.

The two-sided Mann Whitney U test was used to investigate whether there is statistical significance between populations of iNeurons grown in the presence vs. absence of laminin. When running the statistical tests, the images of the different nodes and voltage traces of the different electrodes of a sample were assumed to be independent.

### 2.5 Protocol optimization to enhance survival

#### 2.5.1 Survival rate

To estimate the survival rate, iNeurons were seeded in PDL-coated glass bottom dishes containing two PDMS microstructures each. Two conditions were tested: culturing samples with regular medium (2 samples) and with medium containing 1 μg/mL of laminin (2 samples). 1 h after seeding, the samples were stained with CMFDA and ethidium homodimer-1. The samples were imaged 2 to 4 h after seeding. At DIV 11, the same samples were stained with Calcein AM and ethidium homodimer-1 and imaged. Images of circuits were then cropped into four individual images of nodes (N = 240 nodes per condition).

The number of dead and live cells at DIV 0 was estimated by separately processing the red and green channels of the image of each node. For dead cells, the red channel image was smoothed using a mean filter with a radius of one pixel. Default thresholding was then used to obtain a binary mask ^54^, followed by the built-in Fiji Watershed algorithm to separate neighbouring particles. Finally, the number of particles was counted for each mask(see Fig. †S2a for examples of image processing). Because some of the live cells were slightly overlapping on the green channel images, using this segmentation approach for the live cell count resulted in under-counting the cells. To mitigate the issue of undercounting, a mean filter with a radius of one pixel was applied to the green channel, followed by a minimum filter with a radius of 2 pixels to separate neighbouring cells. Instead of doing segmentation, local maxima were then detected using the built-in Fiji function “Find Maxima”, setting the prominence (maximum height difference between points that are not counted as separate maxima) at 100. The number of maxima detected was used as the live cell count (see Fig. †S2b). The survival rate of node *i* at DIV 0 (*r*_0,*i*_) was calculated for each node as:

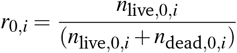

with *n*_*live*,0,*i*_ the number of live cells and *n*_*dead*,0,*i*_ the number of dead cells at DIV 0 for node *i*.

At DIV 11, the number of live cells per node was manually counted. However, many of the dead cells had degraded into several pieces and were overlapping, making it difficult to get a reliable count of dead cells from the red channel images (see Fig. †S2d). We thus made the assumption that the total number of cells per node did not vary between DIV 0 and DIV 11 and used the total cell count from DIV 0 for each respective node to calculate the survival rate. The survival rate of node *i* at DIV 11 (*r*_11,*i*_) was calculated for each node as:

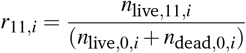

with *n*_live,11,*i*_ the number of live cells at DIV 11 for node *i*.

#### 2.5.2 Area measurement of green-and red-stained structures

##### Staining in open cultures

To investigate survival on open surfaces, iNeurons were seeded in a PDL-coated 48-well glassbottom plate without PDMS microstructures. Two different conditions were tested: culturing the samples with regular medium (9 wells) and with medium containing 1 μg/mL of laminin (9 wells). At DIV 0, 1, 2, and 3, two wells of each condition were stained with CMFDA and ethidium homodimer-1 and 15 to 25 fields of view were taken for each well. At DIV 10, one well of each condition was stained with the live/dead and Hoechst stains and 16 fields of view were imaged. The other 16 wells were restained to investigate the effect of early staining on cell survival.

##### Staining in PDMS microstructures

To study their survival over time, iNeurons were seeded in PDL-coated glass bottom dishes containing two PDMS microstructures each. Two different conditions were tested: culturing samples with medium supplemented with 1 μg/mL of laminin (3 samples) and 10 μg/mL of laminin (3 samples). At DIV 1, 4, and 7, one sample of each condition (each containing two microstructures) was stained with CMFDA and ethidium homodimer-1 and imaged. At DIV 23, the samples imaged at DIV 4 were re-stained with the live-dead and Hoechst stains and imaged. The acquired images were cropped into individual nodes (N = 114 to 120).

##### Area measurement

Due to the difficulty of counting cells in images containing overlapping and degrading cells, the area occupied by green-and red-stained structures was used as an indicator of the evolution of live and dead cells over time. A Gaussian blur filter with a standard deviation of one was applied to each image (red and green channels). For green channel images, Otsu thresholding ^55^ was used and for red channel images, Default thresholding (a Fiji variation of the IsoData algorithm) was used to create a mask from which the area of the objects bigger than 10 pixels was measured. To calculate the percentage of the area occupied by green-or red-stained structures, the measured area (in μm^2^) was either divided by the total area of a field of view (636×636 μm^2^) or by the area of a node (22 700 μm^2^). Images where the resulting percentage was higher than 50 % were visually checked and excluded if the mask obtained did not correspond to the red- or green-stained areas visible on the images.

#### 2.5.3 Protocol optimization

To optimize the survival of iNeurons in PDMS microstructures, several parameters of the protocol were varied: the starting cell density (30 k or 65 k cells/cm^2^), the diameter of the nodes of the microstructure (100 or 170 μm), and the concentration of laminin in the NBD medium (0, 1, or 10 μg/mL). PDL-coated glass bottom dishes with two microstructures each were used as substrates. The samples were stained with live/dead and Hoechst stains at DIV 18 to 23 and imaged. The acquired images were split into individual circuits (N = 30 to 61 circuits per condition). To compare the effect of changing these protocol parameters, the images of individual circuits were visually inspected and the number of nodes within a circuit that had at least one live iNeuron was counted. The number of cells per node was also manually counted.

#### 2.5.4 Axon guidance in microstructures

During the protocol optimization process, many images of circuits with only one node containing live iNeurons were acquired (N = 325 circuits). These were inspected to count the number of nodes with axons growing in the intended, i.e. clockwise, direction and the number of nodes where they did not.

### 2.6 Electrophysiology

#### 2.6.1 Electrical activity recording

The spontaneous electrical activity of 3 MEAs without laminin and 3 MEAs where laminin was added at 1 μg/mL in the cell medium for the first week was recorded for 19 weeks. Recordings were started at DIV 14, performed weekly until DIV 50, and then every other week until DIV 133. During recording sessions, each MEA was taken out of the incubator and placed in the MEA headstage (MEA2100-Systems, Multi Channel Systems), heated to 37 °C with a temperature controller (TCO2, Multi Channel Systems), and kept at 5 % CO_2_ (Pecon #0506.00). The MEA was left in the headstage for 5-10 min to settle before starting the recording. Data was then acquired from the 60 electrodes at 20 kHz for 5 min.

#### 2.6.2 Electrical activity processing

Raw data were band-passed filtered (4th order acausal Butterworth filter, 200-3500 Hz). The baseline noise of the signal was characterized using the median absolute deviation for each electrode for each electrode ^56^. Spikes were detected by identifying negative signal peaks below a threshold of 6 times the baseline noise. Successive events within 1.5 ms were discarded to avoid multiple detection of the same spike. Spike amplitude was defined as the absolute value of the negative amplitude of the detected peak. Spike waveforms were extracted from the filtered voltage trace using the data within a -1ms to 1ms window around the timestamp of the detected spike and used to measure the spike amplitude. Electrophysiological activity was assessed over time by calculating the mean amplitude per electrode, and electrode firing rate, calculated as spike count per electrode divided by the recording time. Mean firing rate per circuit and mean amplitude were both calculated over the active electrodes only. An electrode was considered active if its featured firing rate was above 0.1 Hz.

## 3 Results and Discussion

We report the use of thawed cryopreserved human iNeurons to build biological neuronal circuits composed of less than 50 cells with predominantly unidirectional connections. Circuits are formed using PDMS microstructures consisting of two layers: a first layer with a height of about 200 μm, with through-holes of either 100 or 170 μm of diameter (“nodes”); and a second layer with a height of about 4 μm, connecting the holes through narrow apertures (“microchannels”) (Fig. 1a and b). The PDMS microstructure is placed on a poly-D-lysine (PDL)-coated glass coverslip or microelectrode array (MEA) with channels facing down. PDL is a widely used coating for neuronal cultures due to its good neural adhesion capability, which comes from its positive charge ^57^. After adding cell medium, thawed iNeurons can be seeded by pipetting them on top of the microstructure. iNeurons tend not to adhere to the top of the PDMS but sediment either inside of the nodes or around the PDMS membrane. This is likely due to the hydrophobicity of the PDMS and the flow created when pipetting the medium or moving the sample. Seeded neurons slowly adhere to the PDL-coated surface of the bottom of the nodes. The low height of the microchannels ensures the somas stay in the nodes, preventing cells from migrating out of the nodes into the channels. After a few hours, neurites start extending from the soma and one maturates into an axon, which can grow into the microchannels to connect the nodes together. The channels are designed for axon guidance, leading to mostly clockwise connections between the nodes (see Fig. 6 and Forró *et al*. ^20^). The PDMS microstructures are designed to be placed on top of a 60-electrode MEA, aligning one electrode under each connecting microchannel of a 4-node circuit (Fig. 1c). This allows for recording from the bundle of axons connecting one node to the next.

### 3.1 iNeuron survival over time

#### Survival rate in PDMS microstructures

To check if human iNeurons could form circuits in PDMS microstructures, iNeurons were seeded in PDMS microstructures on PDL-functionalized glass. From their initial spherical shape upon seeding (Fig. 2a) iNeurons spread over the PDL coating and their growing axons formed connections between the nodes of a circuit (Fig. 2b). However, most of the iNeurons died within 2 weeks. The survival rate was quantified by staining iNeurons circuits at DIV 0 and staining these same circuits at DIV 11. With a starting cell number of roughly 70 to 80 neurons per node, the survival rate of iNeurons 4 h after seeding was around 73 % (Fig. 2c). Cell death in the first hours after thawing is likely a consequence of the freezing and thawing process, which puts stress on the cells. Eleven days later, the survival rate in the same circuits dropped to around 0.4 % (Fig. 2d).

**Fig. 2.**
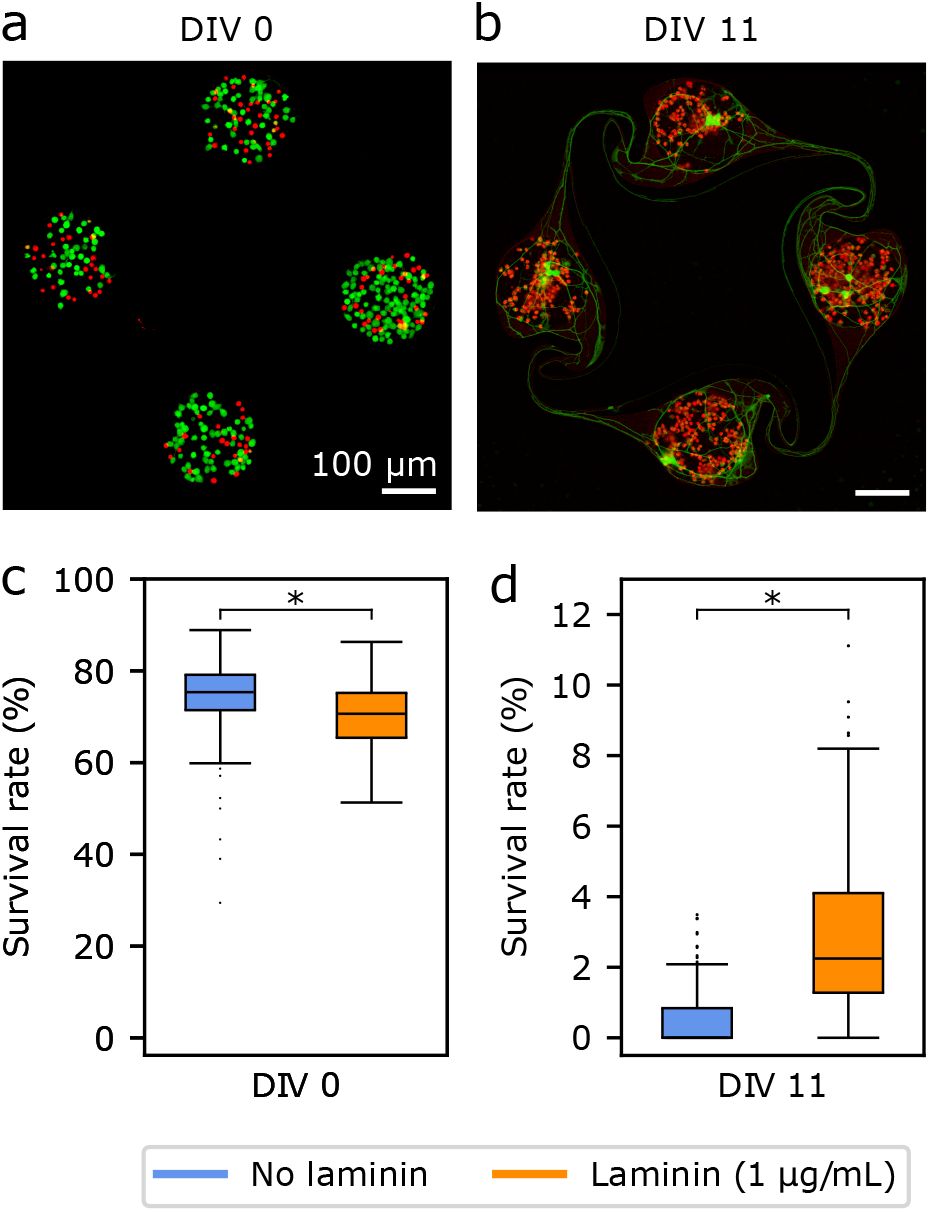
Survival rate over time of iNeurons cultured in PDMS microstructures. (a) Representative example of fluorescently labelled iNeurons grown on a PDL-coated surface at DIV 0 (green: live cells, stained with CMFDA; red: dead cells, stained with ethidium homodimer-1). (b) Same circuit as in (a) at DIV 11 (green: live cells, stained with Calcein AM; red: dead cells, stained with ethidium homodimer-1). For both (a) and (b), the iNeurons were cultured in medium supplemented with 1 μg/mL of laminin. (c) Average survival rate per node for iNeurons cultured in regular medium (blue) and in medium supplemented with 1 μg/mL of laminin (orange), at DIV 0 and (d) DIV 11. For each bar, N = 237 to 239 nodes. *: p < 0.01 (Mann Whitney U test)

Laminin is one of the main components of the basement membrane in the brain and was reported to improve survival and neurite growth in several studies using iNeurons ^43,58,59^. Laminin was thus tested as a secondary coating on top of the PDL. It resulted in a poorer adhesion of the PDMS membrane to the glass and frequent detachment. Adding laminin as a secondary coating after the adhesion of the PDMS membrane to the PDL-coated glass was also unsuccessful. This led to a change of the hydrophobicity of the PDMS, likely due to laminin binding to it. It also caused the iNeurons to stick and grow on top of the PDMS membrane instead of sedimenting to the bottom of the nodes. Finally, we decided to add laminin to the cell medium after seeding the iNeurons into the PDMS microstructure.

Interestingly, in the samples where laminin was added to the medium at a concentration of 1 μg/mL, the survival rate at DIV 0 was of 67 %, which was significantly lower than in the samples without laminin (Fig. 2c). Because the laminin was added at the same time as the CMFDA/ethidium homodimer-1 stains, the difference in the initial survival between the “no laminin” and “laminin” conditions cannot be attributed to the addition of laminin. Instead, that difference is likely due to stochastic variations in the number and survival of cells contained in the volume pipetted onto the sample during the initial cell seeding. The survival rate at DIV 11 in the laminin-supplemented samples was 2.9 % (see Fig. 2d), a significant increase compared to the survival in samples without laminin. Unlike the difference observed in survival rate between both conditions at DIV 0, the increase in survival at DIV 11 can be attributed to the addition of laminin in the medium.

#### Survival of iNeurons in open cultures over time

Supple-menting the culture medium with laminin was beneficial to iNeuron survival, but viability after DIV 11 was still low. Live staining of iNeurons in the early days of culture had a negative impact on the overall cell survival (see Fig. †S3) and should be avoided. In addition to staining, the cause of high mortality in iNeuron circuits might be multi-fold: it could be inherent to thawed iNeurons; to missing factors in the medium; or it could be specifically due to the iNeurons being constrained in small nodes surrounded by PDMS and with comparatively few neighbouring cells compared to *in vivo* conditions. To investigate survival in the absence of PDMS microstructures, iNeurons were plated on bare PDL-coated glass at a high density (300 k cells/cm^2^). Live and dead cells were stained in different samples at different timepoints (DIV 0, 1, 2, 3, and 10), both in regular medium and in 1 μg/mL laminin-supplemented medium (Fig. 3a and b). From DIV 2, live iNeurons tended to cluster and overlap, making it difficult to reliably count the number of cells per field of view (Fig. †S4). For that reason, the area of the field of view occupied by green- or red-stained structures was used as a proxy for investigating the change over time of live and dead neurons. The area occupied by live and dead cells was hypothesized to correlate with the number of live and dead cells.

**Fig. 3.**
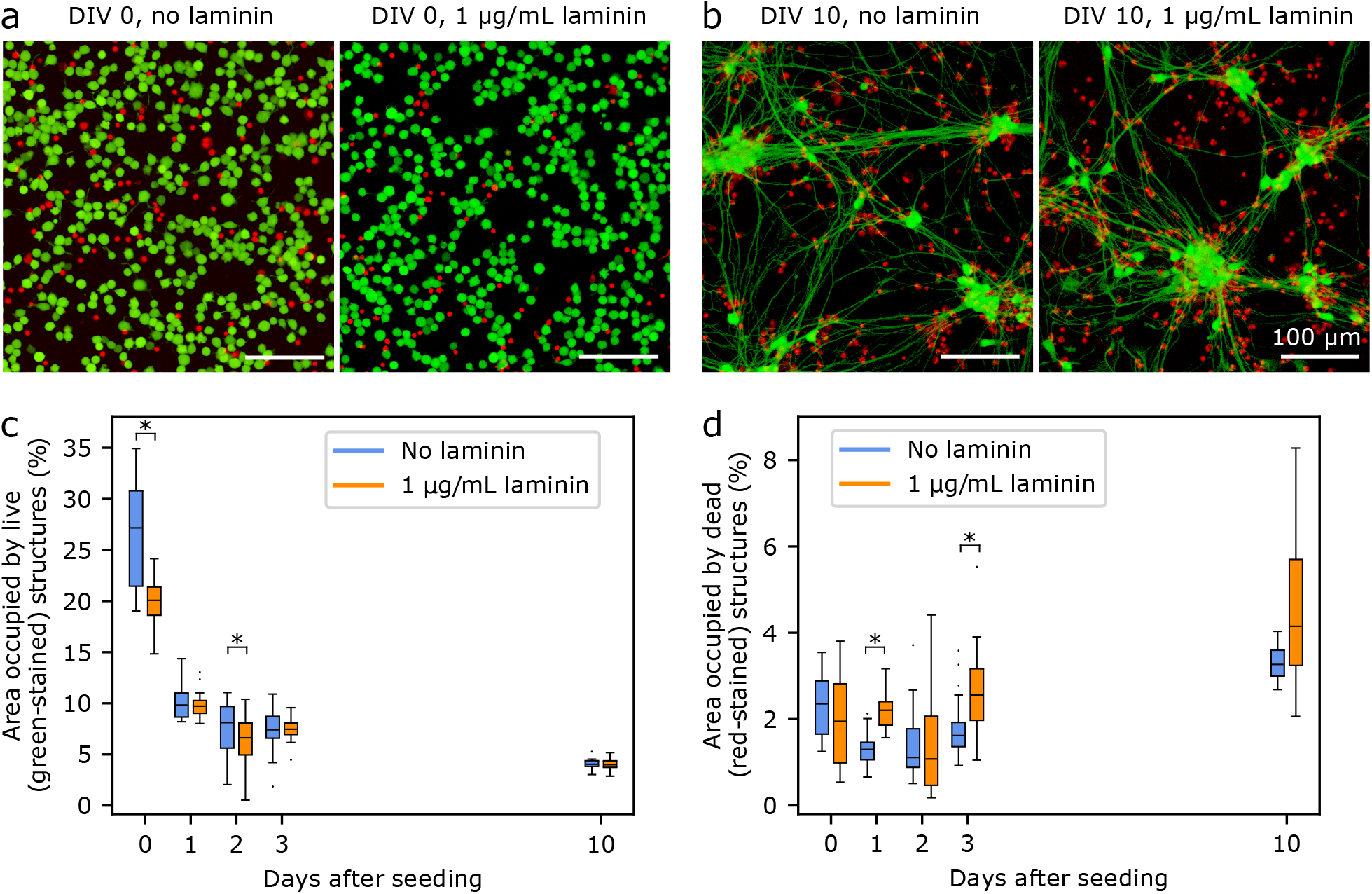
Change over time of the area occupied by live and dead cells in open cultures of iNeurons. (a) Representative example of fluorescently labelled iNeurons grown on a PDL-coated surface at DIV 0, in regular medium (left) and in medium supplemented with 1 μg/mL of laminin (right). (b) Representative example of fluorescently labelled iNeurons grown on a PDL-coated surface at DIV 10 for the same two conditions. For both (a) and (b), live cells are labelled with the green stain Calcein AM and dead cells are labelled with the red stain ethidium homodimer-1. (c) Quantification of the change of average area occupied by green-stained structures (live cells) over time. (d) Quantification of the change of average area occupied by red-stained structures (dead cells) over time. For each box in (c) and (d), N = 13 to 50 fields of view, taken at random in open cultures of iNeurons. *: p < 0.01 (Mann Whitney U test).

The area occupied by live iNeurons decreases by a factor of 4 in the first two days of cultures, before gradually decreasing until DIV 10 (Fig. 3c). Adding laminin to the medium does not have a measurable impact on the area occupied by live cells at DIV 10. There is a statistically significant difference in the area occupied by live cells at DIV 0 between the “no laminin” and “laminin” conditions, but this is again likely due to stochastic differences in the number of cells initially pipetted onto the substrates. Based on the red-stained area measurements, the number of dead cells is fairly constant over the first three days, before slowly increasing between DIV 3 and DIV 10 (Fig. 3d). The area occupied by dead cells is on average higher in the “laminin” than in the “no laminin” condition at DIV 1, 3 and 10 (with a stastistically significant difference at DIV 1 and 3). This could be explained by the fact that laminin tends to make the surface of the substrate slightly more cell-adhesive than bare PDL, leading to more dead cells adhering to it rather than getting washed away during medium changes. Overall, even in an open culture surface at a rather high cell density, many of the iNeurons die over time, especially in the first few days of culture.

#### Survival of iNeurons in PDMS microstructures over time

In order to better understand the evolution of the cell death in circuits over time, iNeurons were cultured in microstructures over three weeks. Since laminin was observed to have a beneficial effect on survival in PDMS microstructure cultures, it was added to the medium in the first week of cell culture, at a concentration of either 1 or 10 μg/mL. We observed that the decrease in the number of live iNeuron per node seemed to take place over the first week of culture, so timepoints for live and dead stains were chosen at DIV 1, 4, 7 and 23 (Fig. 4). At all timepoints, live and dead iNeurons tended to cluster and overlap making it difficult to reliably count the number of cells per node (Fig. †S6). Similar to the open culture results, the area occupied by green- and red-stained structures was used as an indicator of the evolution of the number of live and dead cells over time.

**Fig. 4.**
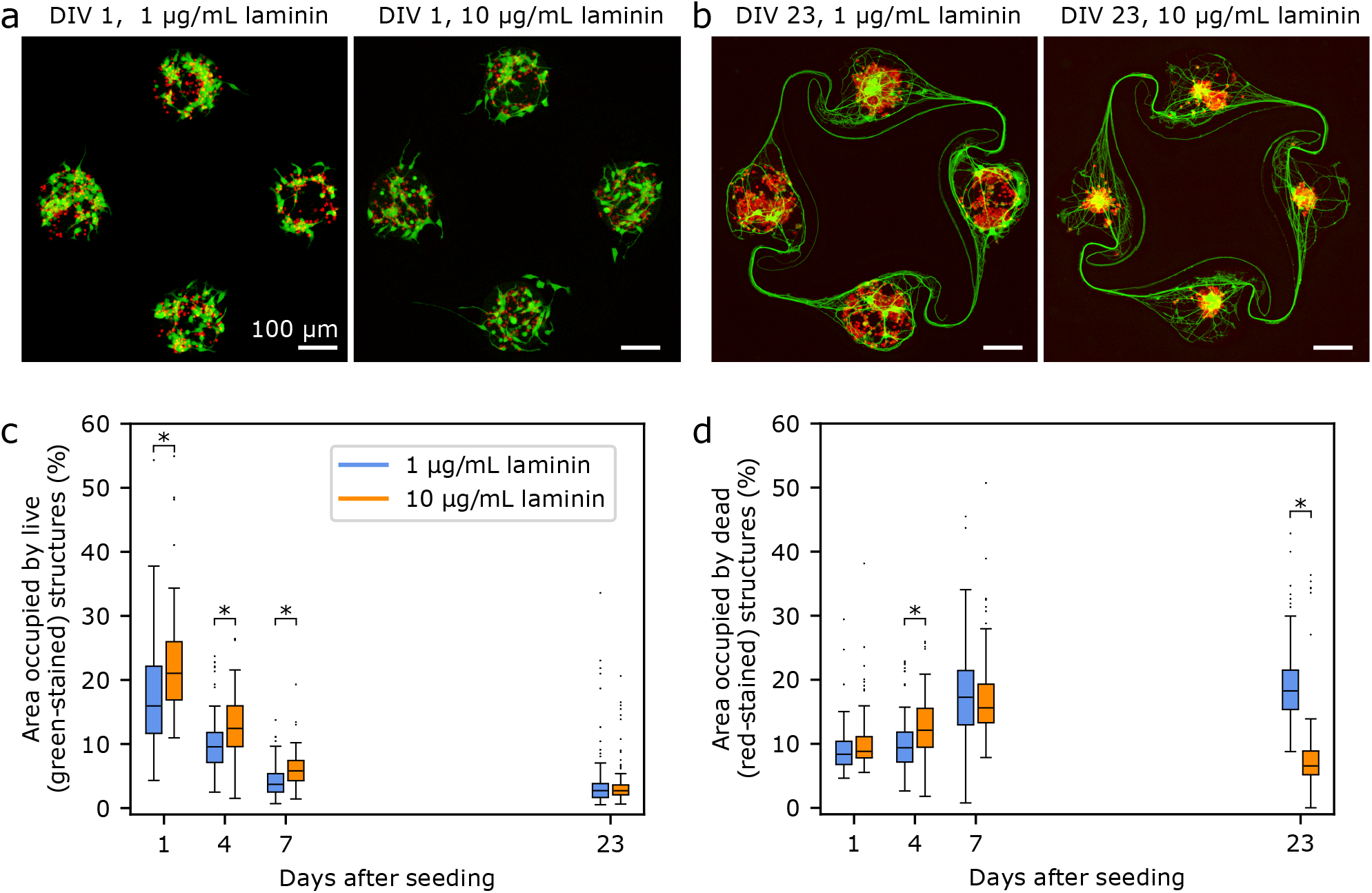
Change over time of the area occupied by live and dead cells for iNeurons cultured in PDMS microstructures. (a) Representative example of fluorescently labelled iNeurons grown on a PDL-coated surface at DIV 1, in medium supplemented with 1 μg/mL (left) and 10 μg/mL of laminin (right). (b) Representative example of fluorescently labelled iNeurons grown on a PDL-coated surface at DIV 23 for the same two conditions. For both (a) and (b), live cells are labelled with the green stain Calcein AM and dead cells are labelled with the red stain ethidium homodimer-1. (c) Quantification of the change of the average area of a node occupied by green-stained structures (live cells) over time. For each point, N = 117 to 120 nodes. (d) Quantification of the change of the average area of a node occupied by red-stained structures (dead cells) over time. For each point, N = 114 to 120 nodes. *: p < 0.01 (Mann Whitney U test).

Based on the area measurement of live structures, the number of live iNeurons in circuits steadily decreases during the first week in culture (Fig. 4c). This is consistent with a qualitative inspection of the images (Fig. 4a and b, Fig. †S6). The area occupied by live cells varied little between DIV 7 and DIV 23, suggesting a stabilization of cell death past a week in culture. The higher (10 μg/mL) laminin concentration lead to significantly higher areas occupied by live cells in the first week of culture. The area occupied by red-stained structures followed an inverse trend to that of the green-stained structures, steadily increasing during the first week in culture before stabilizing. An exception to this is the 10 μg/mL laminin-supplemented samples at DIV 23 where the area occupied by dead structures dropped back to slightly lower levels than on DIV 0. Examining the images, dead cells appear to have clustered under the live cells at DIV 23 (Fig. 4b, right), which could explain the decrease in the area occupied by dead cells. Live cells also seem to cluster together in the center of the nodes more often at the higher laminin concentration. This clustering is due to the fact that live neurons exert forces on each other and on dead neurons, pulling them to the center of the node over time. Higher laminin concentrations might increase the interactions between live and dead cells. In nodes where no iNeurons survived, dead cells did not cluster to the center of the node.

In both cultures of iNeurons on open surfaces and inside of PDMS microstructures the number of live cells drastically reduces over time. Two main differences could be observed between open and PDMS cultures: the time scale in the decrease of live cells and the evolution of the number of dead cells. First, on open surfaces, most of the live area coverage decrease takes place over the first two days in culture, whereas in the PDMS microstructures, the decrease is more gradual and takes place over the first week. Second, on open surfaces, the area occupied by dead cells only slightly increases over time, whereas in PDMS microstructures, it increases inversely to the area occupied by live cells (except for the high laminin sample at DIV 23, as discussed above). When iNeurons are cultured on an open surface, dead cells loosely adhered to the surface likely get detached upon medium change and during the staining steps. Only dead cells that are strongly adhered to the surface will appear on the stained images. In contrast, in the PDMS microstructures, the flow is not strong enough to wash away the dead cells present at the bottom of the cylindrical nodes. As cells die they accumulate inside of the nodes explaining the increase in the number of dead cells that can be observed in Fig. 4d. The accumulation of dead cells in the confined space of a PDMS node might also explain the difference in the time scale of cell death between PDMS microstructures and open cultures: in PDMS microstructures dead neurons diffuse necrotic factors, which can in turn lead to poor survival of the surrounding cells.

In open cultures, we also observed that from around DIV 7-10, iNeurons formed a sheet that tended to easily detach from the surface upon medium changes and staining, requiring extreme care upon handling. This unwanted cell washout was already reported elsewhere ^43^ and limits the possibilities for cell staining past DIV 10. Because iNeurons are not mature at that stage, this can be quite a limitation when performing staining assays on iNeurons. The presence of the PDMS microstructures overcomes this problem, as circuits of iNeurons are protected from turbulent flow by the presence of the PDMS structure.

### 3.2 Optimizing the culture protocol for circuits of iNeurons in microstructures

Because our goal was to record electrophysiological activity from circuits, it was necessary to improve the culturing protocol to obtain better survival of iNeurons in PDMS microstructures. Thawed iNeurons are a fragile cell type and maintaining high viability over the first weeks in culture does not seem to be an attainable goal. However, as the viability stabilizes after approximately 10 DIV, a more pragmatic way to run electrophysiology measurements on circuits is to have at least one live iNeuron per node and a fully connected circuit. We used two metrics to compare the effect of variations in the culturing protocol: first, the number of nodes with at least one live iNeuron; second, the number of live iNeurons per circuit. Both of these were counted in samples cultured for three weeks (DIV 18 to 23). Examples of circuits with 1, 2, 3, or 4 nodes with at least one live iNeuron can be seen in Fig. 5a. The advantage of this metric compared to simply counting the number of live iNeurons per node is that it is less dependent on the starting seeding number of cells. Across a single sample, the initial number of iNeurons per node is expected to follow a Poisson distribution and across different samples, the number of seeded iNeurons depends on the amount of iNeurons present in the volume pipetted during the initial cell seeding, which varies from one sample to the next.

**Fig. 5.**
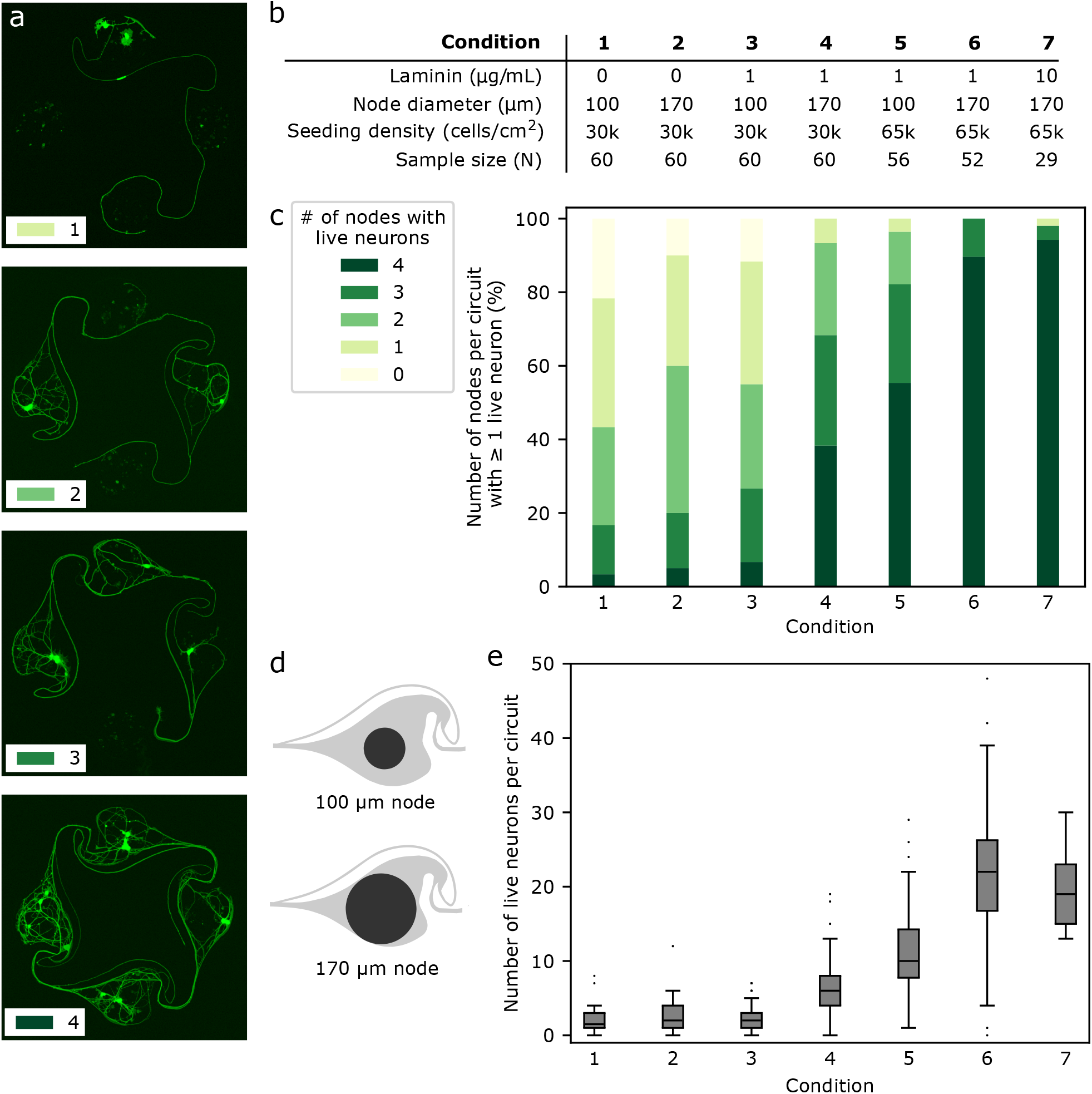
Protocol optimization for the culture of iNeurons in PDMS microstructures. (a) Examples of circuits with 1, 2, 3, and 4 nodes that contain at least one live iNeuron. This metric was used to assess the effect of a parameter change in the cell culture protocol. (b) List of conditions tested to improve the cell survival in the PDMS microstructures. Parameters varied were: supplementing the cell medium with laminin (1-10 μg/mL); increasing the node size (as illustrated in d); increasing the initial cell seeding density from 30 k cells/cm^2^ to 65 k cells/cm^2^. (c) Effect of the different conditions on the percentage of circuits with 0-4 nodes containing at least one live iNeuron at DIV 18 to 23. (d) Two designs were tested: nodes of 100 μm diameter and nodes of 170 μm diameter. (e) Number of live neurons per circuit after three weeks in culture, for all the tested conditions. Mann Whitney U tests were ran on each pair of conditions and the resulting p-values can be found in Table †S1.

Many parameters can contribute to iNeuron death inside of PDMS microstructures: missing factors in the medium; poorly treated PDMS; too low density of neurons; bad nutrient diffusion or too high concentration of necrotic factors. These different parameters were tested to investigate how to positively influence the iNeuron survival in PDMS microstructures.

Despite undeniable advantages such as simple microfabrication and low cost, PDMS can have a deleterious effect on the survival of a neuronal culture, either by slowly releasing uncrosslinked oligomers into the cell culture over time, or by restricting the available nutrients and the removal of waste ^60,61^. Results obtained with open cultures indicated that cell death was high even in the absence of PDMS microstructures (Fig. 3c), but we still tested the effect of cleaning the PDMS microstructures prior to making the substrates. iNeuron survival was compared across non-treated PDMS, ethanol-rinsed PDMS, autoclaved PDMS and extracted PDMS, as described by Millet *et al*. ^61^ These different PDMS treatments did not improve the number of circuits with full nodes (see Fig. †S8), confirming that the release of cytotoxic molecules from the PDMS is not one of the mechanisms behind the low iNeuron survival inside of PDMS microstructures. Another possible cause was the accumulation of dead iNeurons inside of the PDMS node and the poor waste removal possibilities. To test for this, macrophages were added to the iNeuron cultures at DIV 4. While adding macrophages resulted in interesting circuit morphology, it did not seem to affect the percentage of full circuits (see Fig. †S9 and Fig. †S10). Addition of macrophages also complicates the protocol, so this direction was not further explored.

To test if improved nutrient diffusion could help with survival, the diameter of the nodes of the PDMS circuit was increased from the original 100 μm (as designed by Forró *et al*. ^20^) to 170 μm (Fig. 5d). The 170 μm diameter design was already used to obtain the results shown in Figures 2 and 4. As neurons are known to be difficult to culture at low densities and as seeding more iNeurons should equate to more nodes with at least one neuron surviving, the initial cell density was varied from 30 k cells/cm^2^ to 65 k cells/cm^2^. Finally, the effect of adding 1 to 10 μg/mL of laminin to the NBD medium during the first week of culture was tested. Other variations of medium were tested, such as switching from NBD to Neurobasal Plus or BrainPhys, but this did not affect the number of surviving neurons per node (data not shown).

By varying the amount of laminin in the medium, the node diameter and the starting cell density, it was possible to improve the percentage of circuits with at least one live iNeuron per node from 13 % to 94 % (Fig. 5c). Across the seven conditions tested, the median number of iNeurons per circuit after three weeks in culture increased from 2 to 22 (Fig. 5e). To test for the significance of the differences in the number of live iNeurons per circuit, pairwise two-sided Mann Whitney U tests were ran between each pairs of conditions. Detailed p-values for each pair of conditions can be found in Table †S1. Generally, survival in small (100 μm) diameter nodes at a low seeding density (30 k cells/cm^2^) was poor (condition 1). Increasing the node diameter to 170 μm (condition 2) or adding 1 μg/mL of laminin in the cell medium (condition 3) only slightly increased the percentage of full circuits and did not have any significant effect on the average number of live cells per circuit. Combining both (condition 4) resulted in a higher percentage of full circuits and higher average cell number, as did increasing the initial cell seeding density (conditions 5 and 6). Finally, increasing the laminin concentration in the medium could increase the percentage of full circuits to 94 % (condition 7). However, there was no significant difference in the average number of live iNeurons between samples containing laminin concentration of 1 and 10 μg/mL (condition 6 and 7).

Overall, it was possible to optimize the culturing protocol to obtain full circuits with at least one live iNeuron per node in most of the cases, making the protocol suitable to perform electrophysiology recordings on controlled, small circuits of 4 to 50 iNeurons. It is possible to use a higher initial seeding density, but because partial cell death is inevitable, it is not desirable to use too high of a density to avoid clogging the nodes with dead cells.

### 3.3 Axon guidance in microstructures

The initial stomach design with 100 μm diameter nodes was reported by Forró *et al*. to lead to 92 % axon guidance success when seeding rat primary hippocampal neurons. This high percentage of success can be explained by the shape of the chamber: an axon growing towards the counter-clockwise node should get redirected by the curved side channel (see Fig. †S13 and Fig. †S14). To test if axon guidance was also successful with human iNeurons using 170 μm diameter nodes, images of circuits which had only one node containing live iNeurons were inspected. More than 300 such images were obtained during the protocol optimization phase. For 100 μm diameter nodes, the guidance success rate was 90.1 % (N = 223) and for 170 μm diameter nodes, the success rate was 91.2 % (N = 102). Examples of successful axon guidance in 170 μm and 100 μm diameter nodes can be seen in Fig. 6a and b. Examples of unsuccessful axon guidance can be seen in Fig. 6c. These results confirm that the stomach design can be used to get mostly unidirectional connections between the nodes of a circuit containing iNeurons.

**Fig. 6.**
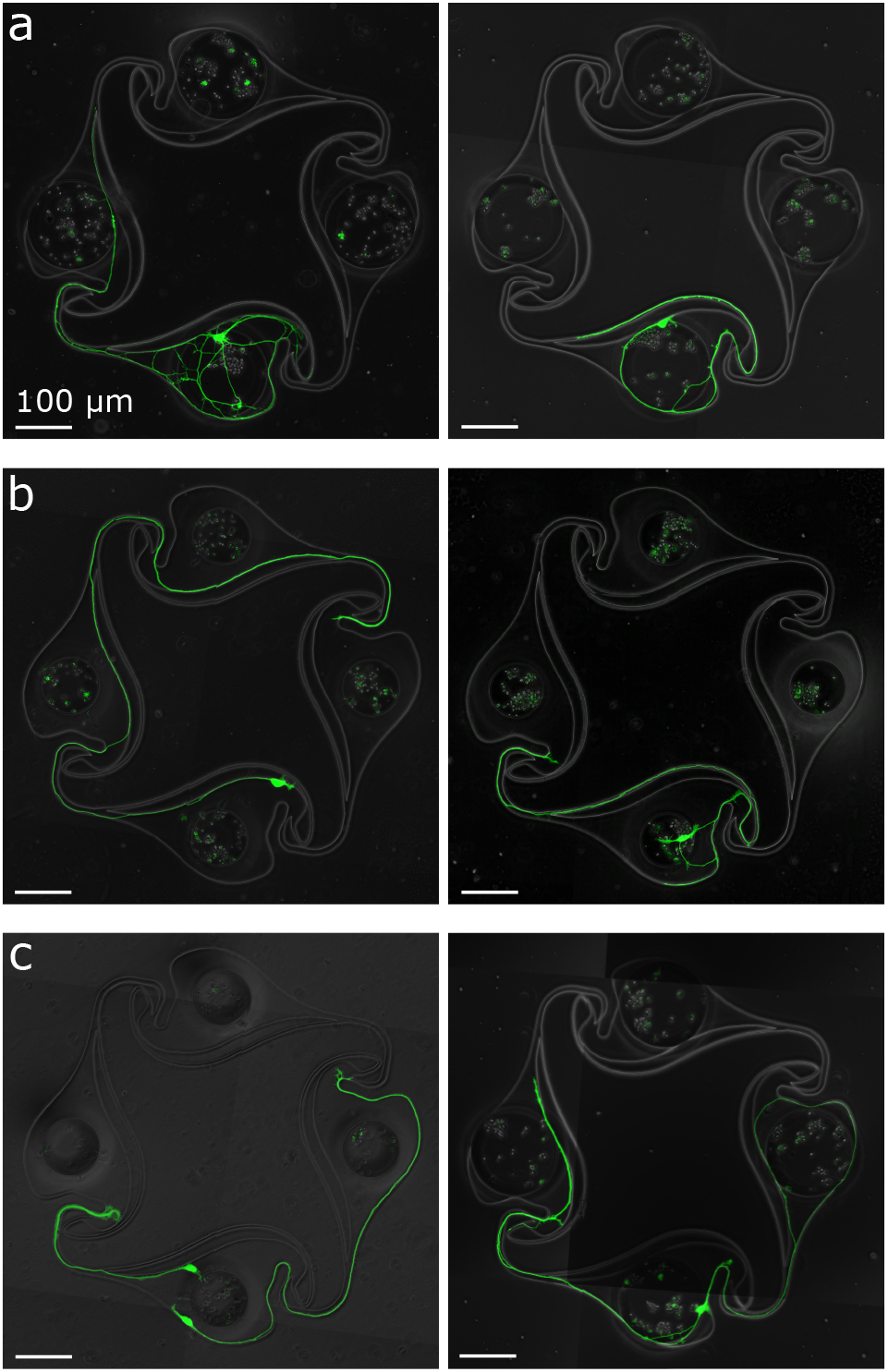
Axon guidance in “stomach” PDMS microstructures, assessed by inspecting circuits with only one node containing live iNeurons. (a) Examples of circuits with 170 μm diameter nodes where the PDMS microstructure successfully guided an axon into the intended clockwise direction. This was the case for 90.1 % of the inspected circuits (N = 223). (b) Same for 100 μm diameter nodes. Success rate was 91.2 % (N = 102). For both (a) and (b), the right-hand side pictures show examples of the successful redirection of an axon into the curved side channel, a particularity of the stomach structure. (c) Examples of circuits where the PDMS microstructures failed to guide the axon into the expected clockwise direction. This was the case for 9.9 % of the inspected circuits (N = 325). In these cases, axons grew towards the counter-clockwise node rather than getting redirected into the side channel. All images displayed here are an overlay of a phase-contrast picture and a fluorescent Calcein AM staining of the iNeurons.

### 3.4 Electrophysiological recordings

Using a protocol optimized for survival of iNeurons in PDMS microstructures, we built arrays of 15 four-node circuits of iNeurons on MEAs and recorded their spontaneous electrical activity across 133 DIV. Fig. 7a and b show an example of a circuit of iNeurons at DIV 35 and DIV 138. This circuit was part of a sample supplemented with 1 μg/mL of laminin during the first week of culture. Examples of raw voltage traces recorded from this circuit can be seen on Fig. 7d. These were recorded from the top left electrode of the circuit (red electrode on Fig. 7c) at different time points (DIV 21, 62, 90, and 133). The action potentials detected from a 5 min recording of spontaneous electrical activity were extracted from the filtered voltage traces and overlaid on Fig. 7e. Overlay of the action potentials detected on the other three electrodes can be seen on Fig. †S15. Raster plots of 40 s of spontaneous electrical activity for this circuit can be seen on Fig. 7f.

**Fig. 7.**
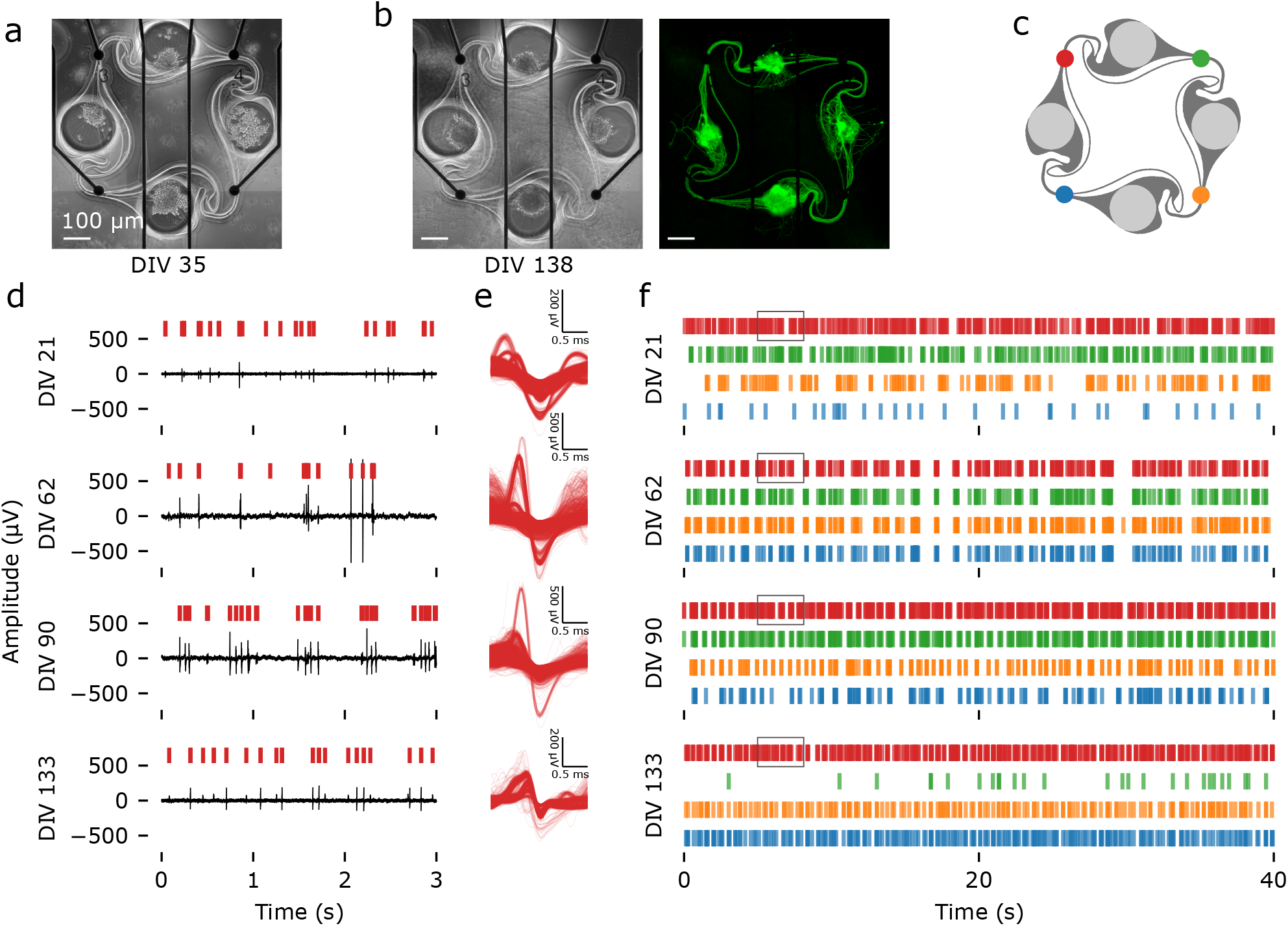
Spontaneous electrical activity over 133 DIV, recorded from an example iNeuron circuit. (a) Images of a circuit of iNeurons aligned to the four electrodes of a MEA at DIV 35 (phase contrast). (b) Same circuit at DIV 138 (left: phase contrast; right: Calcein AM). The iNeurons composing the circuit are still alive and firing after more than four months in culture. (c) Color code for the four electrodes of the circuit. (d) Example of raw voltage traces recorded at the red electrode of the circuit shown in (a) and (b) at four time points (DIV 21, 62, 90 and 133). The red vertical bars indicate detected spikes. (e) Overlay of the waveforms of the action potentials detected in a 5 min recording of spontaneous electrical activity for the red electrode at the same four time points. (f) Raster plot showing the timestamps of the spikes detected during 50 s of recording of the spontaneous electrical activity of the same circuit at four time points. The four colors correspond to the electrodes shown in (c). The grey boxes indicate the time frame corresponding to the raw data showed in (d).

As the microelectrodes are placed below the microchannels where the axons connecting two nodes are growing, the recorded voltage trace consists of spikes coming from the axons rather than from the soma. In open cultures, axonal spikes are small, usually below the noise level, and thus cannot be recorded using MEAs ^62^. In axon-containing PDMS microchannels, axonal spikes are amplified by the presence of the PDMS ^63,64^. This allows to record spikes with high signal-to-noise-ratio, as shown on Fig. 7d and e. The percentage of active electrodes, mean firing rate and mean amplitude over time were calculated from the data recorded from three different 60-electrode MEAs (180 electrodes) with and without the addition of 1 μg/mL of laminin in the medium in the first week of culture. An electrode was considered “active” if its mean firing rate was greater than 0.1 Hz during the weekly 5 min recording of spontaneous activity. At DIV 119, one of the samples without laminin had no active electrodes anymore, likely because all of the iNeurons died. Measurements were thus stopped at DIV 119 for these samples. The samples with laminin still had active electrodes, but recordings were stopped at DIV 133 due to the university closing down for the winter break.

The laminin samples had a higher percentage of active electrodes (Fig. 8a), which can likely be explained by the fact that more cells survived than in the samples without laminin. Past DIV 90, a drop in the percentage of active electrodes is visible in the samples with laminin. This is because around that time, we observed that axons had grown on the upper surface of the PDMS microstructure and performed a live cell staining to further investigate that (Fig. †S16). There were no cell bodies on top of the PDMS, but axons seemed to have grown from the nodes onto the top of the PDMS. The staining likely had an adverse effect on some of the surviving cells. For both conditions, the mean firing rate increased over the first two months, up to DIV 62, before decreasing (Fig. 8b). During that time frame, the mean amplitude regularly increased in the samples with laminin before plateauing (Fig. 8c). The mean amplitude of samples without laminin also increased, but in a less regular manner. There was no statistically significant difference in the mean amplitude between the “laminin” and the “no laminin” samples at any of the time point. For the mean firing rate, the only statistically significant difference that could be observed between both types of samples was at DIV 119 (p < 0.01, two-sided Mann Whitney U test).

**Fig. 8.**
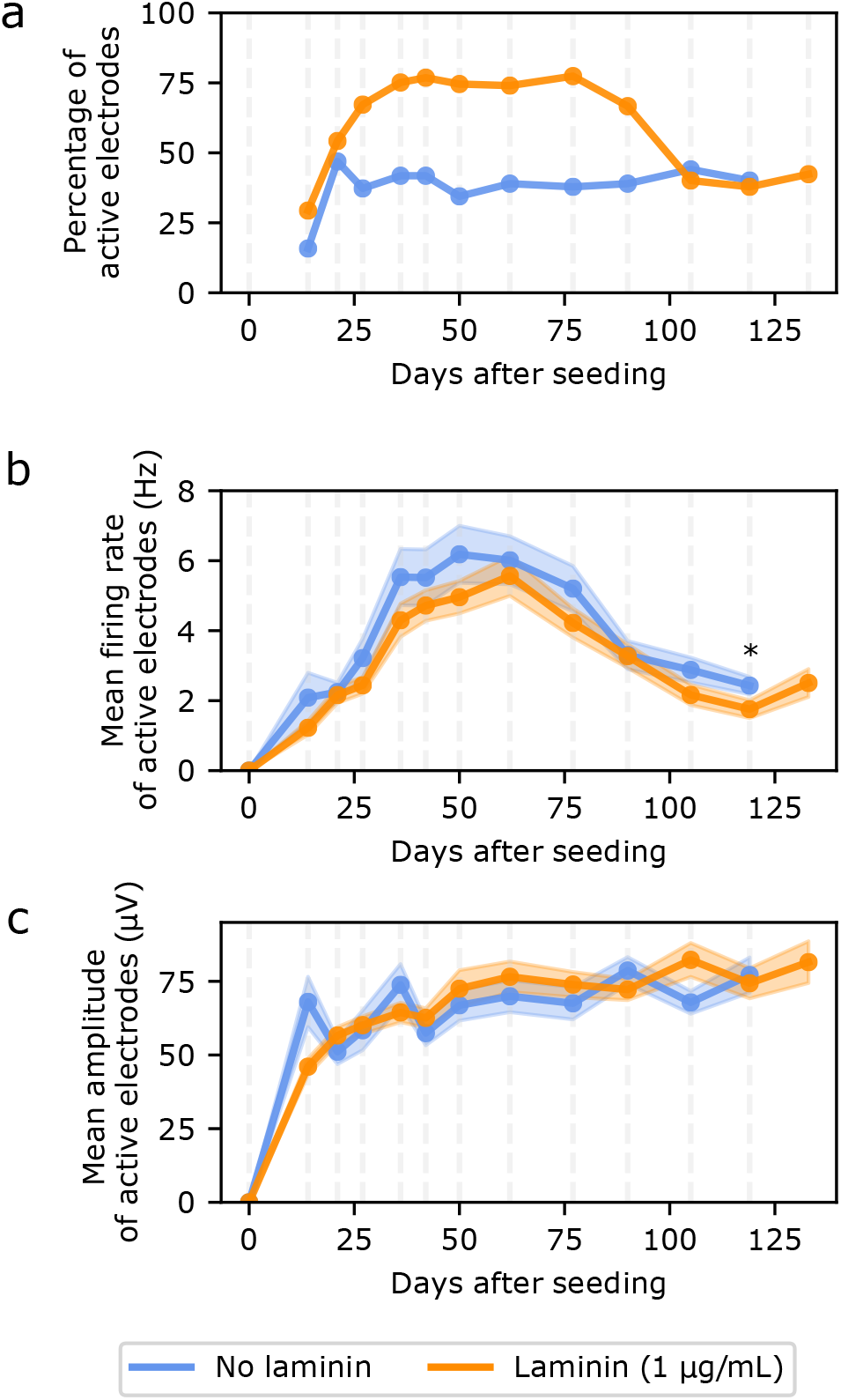
Characterising the spontaneous electrical activity of iNeurons circuits cultured in regular medium (blue) and in medium supplemented with 1 μg/mL of laminin (orange). Data were recorded from 3 MEAs for each condition, at DIV 0, 14, 21, 27, 36, 42, 50, 62, 77, 90, 105, 119, and 133 (“laminin” samples only on DIV 133). (a) Percentage of active electrodes out of 180 electrodes (3 MEAs) for each condition. An electrode was considered active if its firing rate was of at least 0.1 Hz. (b) Mean firing rate of the active electrodes. (c) Mean amplitude of active electrodes. For both (b) and (c), the shaded area represents the SEM and N = 28 to 137 electrodes for each point (corresponding to the percentage of electrodes showed on (a)). *: p < 0.01 (Mann Whitney U test)

Overall, we were able to record electrophysiology data over more than four and a half months, a longer experimental period than what is typically reported in studies using iNeurons ^65,66^. This makes our platform suitable for long-term experiments. leaving enough time for iNeurons to become functionally mature. However, axons start growing on top of the PDMS microstructures after a few weeks, leading to connections between circuits, which is not desirable in regards to keeping independent circuits. This can be overcome by coating the top of the PDMS with an antifouling molecule, such as PAcrAm-g-(PMOXA, amine, silane) ^67^.

## 4 Conclusions

We demonstrated the successful building of circuits of less than 50 thawed cryopreserved human iPSC-derived neurons in PDMS microstructures, some of which can survive over several months. Such a platform can be used for both imaging and long-term electrophysiological assays. To our knowledge, this is the first report of building *in vitro* circuits using human-derived cortical neurons with control over the topology and so few neurons per circuit. Survival of thawed iNeurons is low, but we optimized the protocol to obtain full circuits in most cases.

This technology holds the potential to study fundamental signal processing in neurons. It can, for example, be used to study plasticity by recording the response of a circuit upon stimulation of its electrodes. Because the MEA provides a functional readout of several circuits in parallel, the platform could also potentially be adapted for translational research applications such as testing the effect of neuromodulatory molecules on the electrical activity of a circuit. This might be especially interesting in combination with patient-derived cells with neurological disorders ^65,68,69^ and in particular to study circuit or neurodevelopmental disorders such as Alzheimer’s disease and autism or Fragile X syndrome. We demonstrate that iNeurons can be cultured and electrically probed for several months, leaving them sufficient time to reach maturity levels that could biologically replicate diseases.

Several improvements can be implemented in this system. First, the layout presented here consists in four-node circuits, but could be adapted to answer scientific questions of interest, as needed by the experimenter. Second, the current layout of the PDMS circuit is constrained by the need to interface it with a 60-electrode MEA layout and only allows recording and stimulating from specific positions in the system. By using a high-density CMOS MEA ^70^, such design constraints could be eliminated and any part of the circuits could be recorded from and stimulated. This would however complicate imaging assays, as high-density CMOS MEAs are not transparent. Thirdly, only one cell type was used in this work, but more complex circuits could be built by seeding different types of cells in different nodes of a circuit, for example excitatory and inhibitory neurons to test for spike-timing-dependent plasticity. Such cell types are now commercially available as cryopreserved cells, thanks to recently developed protocols to differentiate iPSCs into brain cell types such as dopaminergic neurons, GABAergic neurons and astrocytes. Seeding different nodes of a circuit with different cell types would require a fine control over cell placement, which can for example be achieved using technologies such as pick-and-place with a modified atomic force microscope (FluidFM) ^71^. Overall, the possibility to build small circuits of human-derived cells that survive over several months holds great promise for advancing fundamental neuroscience research and may also find translational applications.

## Author Contributions

SG and JV designed the research project. SG wrote the manuscript. SJI, SW, JM and CF performed preliminary experiments. SG conducted the experiments with support from JBP. SG analysed the data, with the help of BC for the electrophysiology data. JV secured funding for the projects. IF and MM produced the iPSCs and iPSC-derived neurons. All co-authors reviewed and approved the manuscript.

## Acknowledgements

Warm thanks to Margarita Dinamarca Ceballos from the Pecho-Vrieseling lab for useful advice on culturing iPSC-derived neurons. ETH Zurich, the Swiss National Science Foundation, the Swiss Data Science Center, the FreeNovation grant, the Human Frontiers for Science Program and the OPO Foundation are acknowledged for financial support.

