## Supplementary Information for "Topologically controlled circuits of human iPSC-derived neurons for electrophysiology recordings"

### 1 Image analysis for survival rate estimations

#### 1.1 Survival rate at DIV 0 and DIV 11

Fig. S1 shows representative images of a fluorescently labelled circuit of iNeurons. Such images were used to obtain the plots shown in Fig. 2. The number of live and dead cells on DIV 0 images were analysed according to the method presented in section 2.5.1. of the main paper. Fig. S2a and b show a step-by-step example of the automatized counting of live and dead cells for an example node for both the "no laminin" and "laminin" conditions. Fig. S2c compares the numbers obtained from the images shown in (a) and (b) with the manual hand count. Automatized counting is within 15 % of the manual count. Fig. S2d shows the manual count of the live cells at DIV 11. By DIV 11, dead cells were not uniformly stained and some of them had separated into several pieces, making the use of segmentation unreliable to count the number of dead cells. For that reason, the survival rate at DIV 11 was computed as the number of cells manually counted at DIV 11, divided by the total number of cells automatically counted at DIV 0. This was possible because the same circuits were stained at DIV 0 and restained at DIV 11.

#### 1.2 Effect of CMFDA/Ethidium homodimer-1 staining in early days of culture

We observed that staining cultures at early DIV has an adverse effect on the cell survival at later DIVs. We could quantify that effect by restaining at DIV 10 the wells of the 48-well plate used to obtain the plots shown on Fig. 3. Compared to wells that were stained for the first time at DIV 10, restaining wells that had already been stained at DIV 0, 1, 2 or 3 resulted in 1 to 1.5 times lower area occupied by live cells. Early stains thus seem to have an adverse effect on survival and should be avoided. They can be replaced with genetically expressed fluorescent proteins. However, if using a typical cytosolic fluorescent proteins such as GFP, dead cells might also have expressed the fluorescent protein before dying, leading to unclean fluorescent images that usually cannot be automatically segmented and analyzed.

#### 1.3 Area measurement of green- and red-stained structures

Fig. S4 shows representative images of fluorescently labelled iNeurons at DIV 1, 2 and 3. Such images are the ones that were used to obtain the plots shown on Fig. 3. The area occupied by green- and red-stained structures at the different DIVs was measured according to the method presented in section 2.5.2. of the main text. Fig. S5 shows examples of the steps used to obtain the binary masks from which the area occupied by live and dead cells was calculated. Due to the microscopy settings used, fields of view from DIV 0 were smaller than images from the other DIVs (424 $\mu$m vs 626 $\mu$m of side). This was taken into account by calculating the percentage of the field of view occupied by live- or red-stained structures.

Fig. S6 shows representative images of fluorescent labelled circuits of iNeurons at DIV 4 and 7. Such images are the ones that were used to obtain the plots shown on Fig. 4. Each image of a full circuit was cropped into its four nodes. The area of the node occupied by green- and red-stained structures at the different DIVs was measured according to the method presented in section 2.5.2. of the main text. Fig. S7 shows examples of the steps used to obtain the binary masks from which the area occupied by live and dead cells in the node was calculated. The percentage area was obtained by dividing the white area by the area of the node (2.27x10$^{-}$2 mm$^2$).

### 2 Protocol optimization

#### 2.1 PDMS treatment

To determine if part of the cell death could be attributed to poorly treated PDMS, the effect of different PDMS cleaning methods prior to substrate making was assessed. The cleaning methods tested were autoclaving, solvent extraction and ethanol (A15, Thommen-Furler AG). Autoclaving consisted in placing the PDMS micostructures in an autoclave (Varioklav

75T, Sterico) and heating them at 121°C and 110 kPa for 20 min, followed by a 20 min drying cycle at a temperature of 81°C to 91°C. Solvent extraction was performed according to the extraction protocol reported by Millet *et al.*[1]. Ethanol cleaning consisted in immersing the PDMS membrane in ethanol for approximately 16 h, followed by 24 h of drying in an oven at 60°C.

Fig. S8 shows the distribution of the number of nodes with at least one live iNeuron for the different PDMS cleaning methods. It seems like the different cleaning methods had either little effect (autoclaving) or an adverse effect on the number of full circuits (solvent extraction and ethanol), so none of these PDMS treatments were kept in the substrate preparation protocol.

#### 2.2 Macrophage co-culture

Because of the poor survival rate, dead iNeurons accumulate in the nodes of the PDMS circuits and might have an adverse effect on the remaining live iNeurons in the node. To test for this, iPSC-derived macrophages were added to the cultures on DIV 4. iPSC-derived macrophages were obtained following the protocol described by Giorgetti *et al.*[2] and kindly provided by Novartis as a suspension.

Upon reception, macrophages were centrifuged for 5 min at 1000 rpm, resuspended in macrophage medium and plated into a non-coated 6-well plate (92006, TPP). Macrophage medium consisted in RPMI 1640 GlutaMax (61870-036, ThermoFisher) with 10% heat inactivated FBS (A156-152, ThermoFisher), 1% sodium pyruvate (11360-039, ThermoFisher), 1% pen-strep (15140, ThermoFisher), 50 $\mu$M of mercaptoethanol (31350-010, ThermoFisher) and 40 ng/mL of human M-CSF (216-MC-025, Biotechne). Macrophages were kept in incubator for 8 days. They were then detached using TrypLE (12604-013) and centrifuged at 500 rpm for 5 min. Macrophages were seeded on top of a DIV 4 culture of iNeurons in PDMS microstructures at a density of about 50 k cells/cm$^2$. Phase contrast images of the wells were taken every other day until DIV 20. At DIV 20, a live/dead and Hoechst stain was performed on them.

Fig. S9a shows an example of one node getting cleaned by the macrophages over time. Impressively, the macrophages seem to phagocytose all of the dead cells contained in the nodes. Macrophages can squeeze into the low part of the chamber (see DIV 8 image) and move around the chamber a lot.

Fig. S10 shows a comparison between macrophage-containing and no macrophage-containing cultures. In the presence of macrophages, live iNeurons tend to tightly cluster together and their axons are grouped into fairly straight bundle connecting all four nodes (see Fig. S9b and Fig. S10b). This is likely due to macrophages moving around the nodes and leading to a mechanical bundling of the axons along the most straight path connecting one node to the next. However, the presence of macrophages does not seem to lead to a big improvement on the number of full nodes. Adding macrophages lead to a more complex protocol, along with adding uncertainties of mixing several cell types whose *in vivo* interactions are not fully understood. For those reasons, co-culture of macrophages and iNeurons was not further explored.

#### 2.3 Full circuits

Fig. S11 shows representative examples of circuits with four nodes containing at least one live iNeuron for all of the conditions listed in Fig. 5b. A stitched image of all of the 15 circuits from one PDMS membrane can be seen on Fig. S12. This was obtained with condition 7 (10$\mu$g/mL of laminin).

#### 2.4 Statistical tests on protocol variations

The number of live iNeurons per circuit after 3 weeks (DIV 18 to 23) was counted for all of the conditions presented in Fig. 5b. The statistical significance of the difference of the number of cells per circuit was tested by running pairwise two-sided Mann Whitney U tests on each pair of conditions. Results can be seen in Table S1.

**Table S 1** P-values obtained by running the Mann Whitney U test for all pairs of conditions listed in Fig. 5b. *: $p < 0.01$

| Condition | 2 | 3 | 4 | 5 | 6 | 7 |
|---|---|---|---|---|---|---|
| 1 | 0.10 | 0.02 | 2.14E-13 (*) | 1.60E-18 (*) | 1.63E-19 (*) | 8.84E-15 (*) |
| 2 | | 0.21 | 5.96E-11 (*) | 2.84E-17 (*) | 3.09E-19 (*) | 8.40E-15 (*) |
| 3 | | | 1.12E-10 (*) | 2.75E-17 (*) | 3.88E-19 (*) | 9.44E-15 (*) |
| 4 | | | | 2.01E-07 (*) | 3.35E-17 (*) | 9.79E-14 (*) |
| 5 | | | | | 1.07E-11 (*) | 1.07E-08 (*) |
| 6 | | | | | | 0.13 |

### 3 Directionality

The "stomach" design of the PDMS microstructure allows to guide axons in a clockwise directions in 90% of the cases. This is due to the shape of the chamber and to the properties of the axons. When an axon starts growing from a single soma (Fig S13a), it grows until it hits a wall, then tends to follow the wall. If it grows towards the clockwise direction, it can simply follow the output channel to the next node (Fig S13b and c). If the axon grows towards the counter-clockwise direction, it will in most cases be redirected to the side channel, either because the axon is already following the top wall and naturally continues in the side channel (Fig S13d) or because it cannot follow the sharp angle from the input channel (Fig S13e). In 10% of the cases, the axon manages to follow the sharp angle from the input channel (Fig S13f) and connects in the wrong direction.

All of these different possible cases were observed in circuits where few neurons survived. Examples for each case are shown in Fig S14, for both the 100-$\mu$m (top) and 170-$\mu$m (bottom) diameter node design. Fig S14f shows examples where several neurons were growing in the node.

### 4 Electrophysiology

#### 4.1 Action potential waveforms

The action potential waveforms for all four electrodes of the circuit shown on Fig. 7 were extracted at DIV 36 and DIV 133. They are visible on Fig. S15.

#### 4.2 Axon growth on top of PDMS microstructures

MEA recordings were taken over 133 DIV. Around DIV 90, a CMFDA staining was performed to investigate axons growing on top of the PDMS microstructures. An example of the growth of axons for one of the samples is visible on Fig. S16. On the one hand, almost no live soma are visible on the top surface of the PDMS, consistent with the observations that neurons generally could not survive on top of the PDMS on the first day of seeding. On the other hand, a lot of dead cells are visible on the phase contrast image, seemingly interacting with the network of axons. Over time, axons seem to have grown from the live cells inside of the nodes onto the top surface of the PDMS. This is probably due to the presence of laminin and other proteins in the cell medium, which deposit over time on the surface of the PDMS. Growth of axons on top of the PDMS could be avoided by functionalizing the top surface with an anti-fouling molecule.


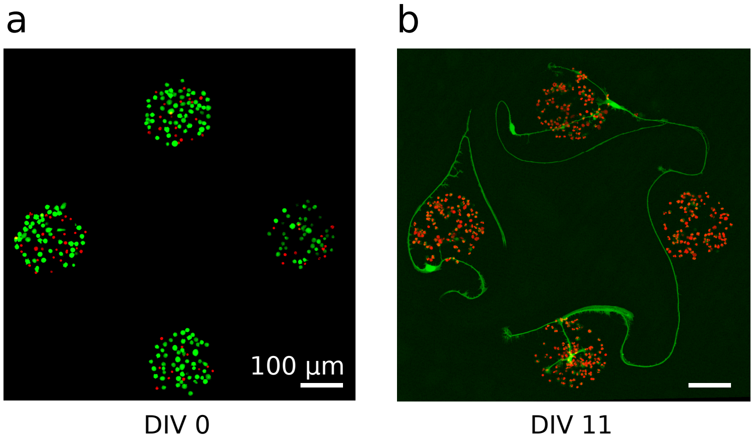


**Figure S 1** Example of a circuit of fluorescently labelled iNeurons grown on a PDL-coated surface at DIV 0 (a) and at DIV 11 (b), without laminin supplemented to the culture medium. Both images come from the same circuit. At DIV 0, the green dye is CMFDA and at DIV 11, the green dye is Calcein AM (both staining live cells). For both, the red dye is ethidium homodimer-1, staining dead cells.

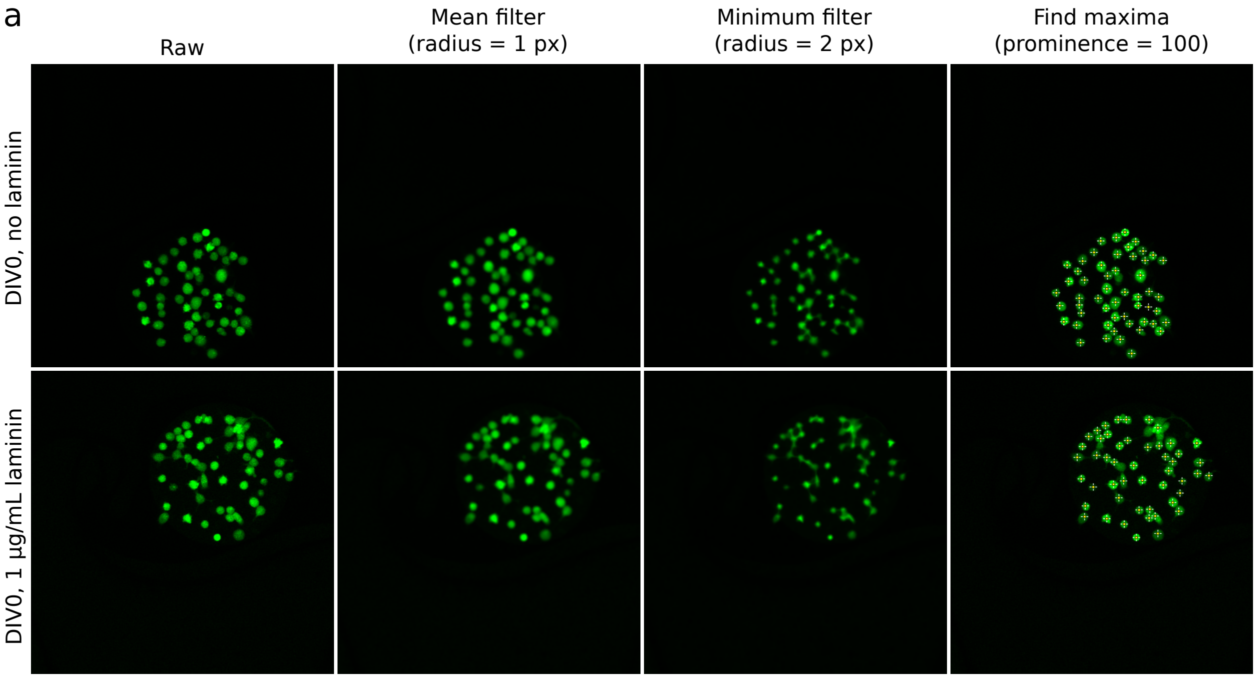


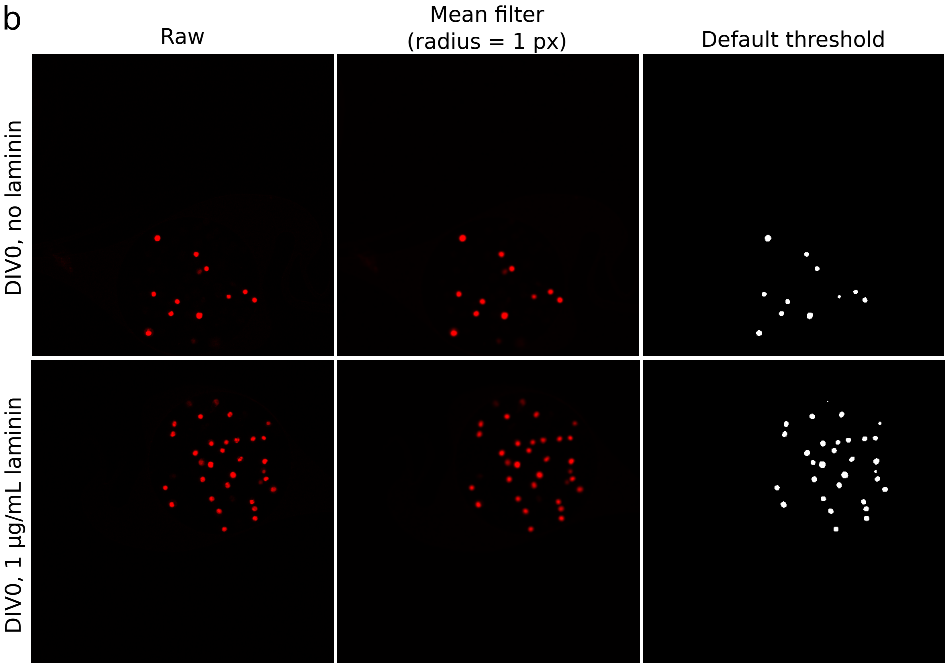


c

| DIV 0 | | Auto count | Hand count |
|---|---|---|---|
| Live | No lam | 11 | 11 |
| | Laminin | 31 | 32 |
| Dead | No lam | 53 | 53 |
| | Laminin | 49 | 56 |

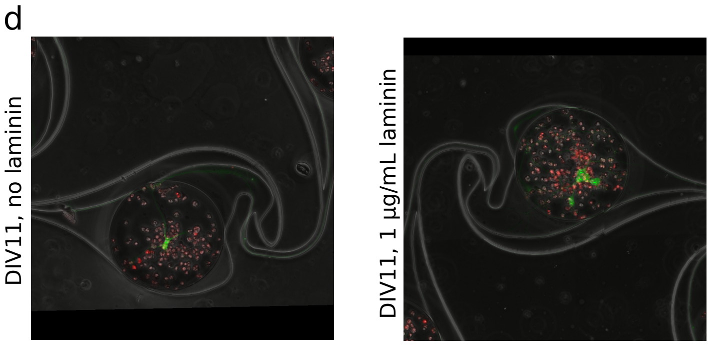


**Figure S 2** Step-by-step image analysis to count the number of live and dead cells in DIV 0 and DIV 11 node images (for both the "no laminin" and the "laminin" conditions). (a) Analysis of nodes containing live cells. px: pixels. (b) Analysis of nodes containing dead cells. (c) Comparison between automated counting of the number of cells per node (using the pipelines shown in a and b) and a handcount of the number of cells per node. (d) At DIV 11, the live cells overlapped and the dead cells were not stained clearly enough to be automatically counted. For that reason, the number of live cell was manually counted, node by node.

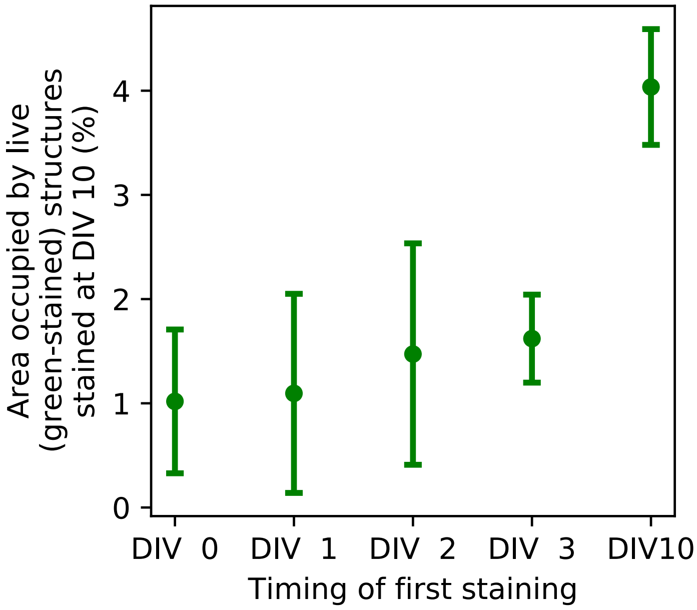


**Figure S 3** Performing a first CMFDA/ethidium homodimer-1 staining in the early days of cell culture decreases the area occupied by green (live) structures at DIV 10, compared to a sample that was stained at DIV 10 for the first time. Staining at DIV 10 consisted in Calcein AM.

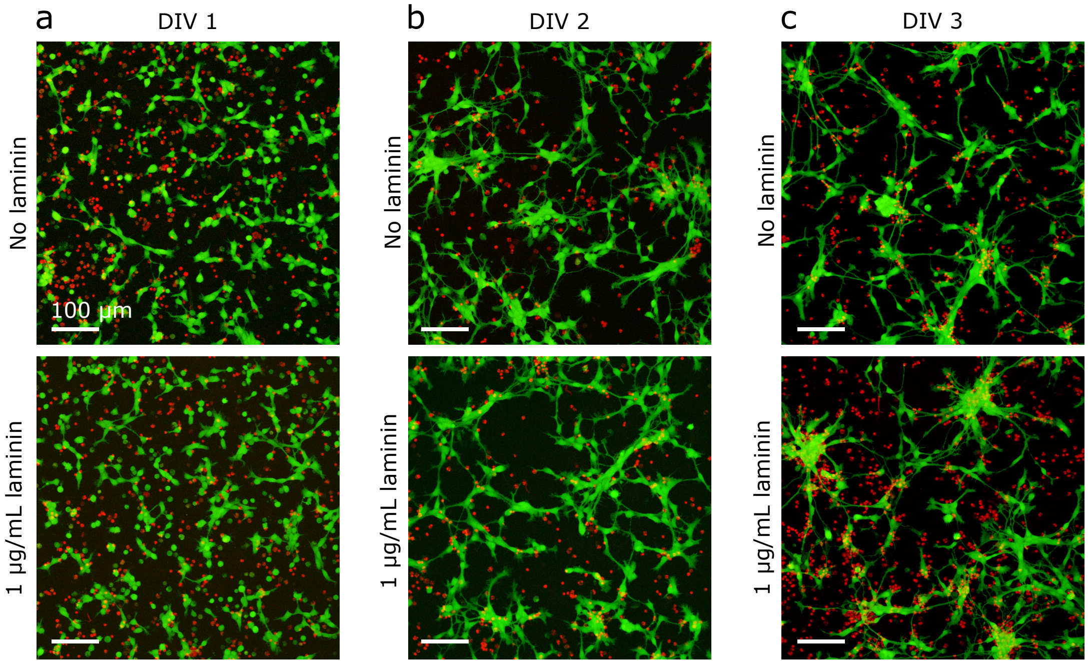


**Figure S 4** Representative examples of fluorescently labelled iNeurons grown on a PDL-coated surface at DIV 1 (a), 2 (b) and 3 (c). Green: live cells, CMFDA. Red: dead cells, ethidium homodimer-1.

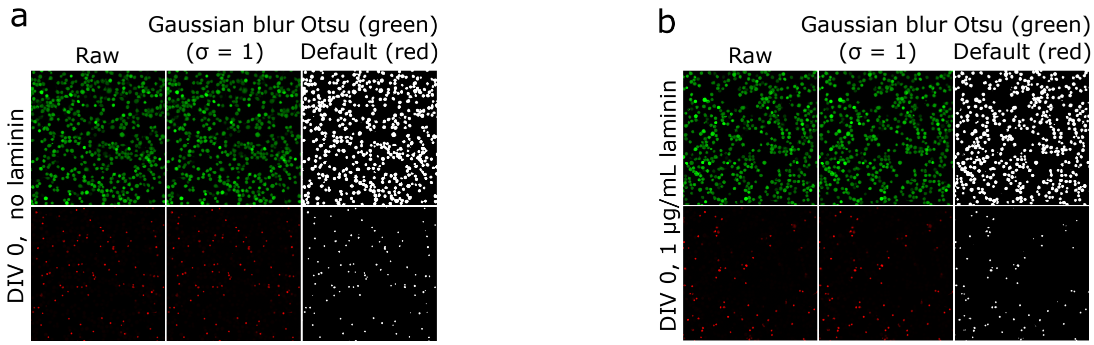


**Figure S 5** [Continued on next page] Step-by-step image analysis to measure the area occupied by live and dead cells in images of open neuron cultures. (a) DIV 0 - no laminin (b) DIV 0 - laminin (c) DIV 1 - no laminin (d) DIV 1 - laminin (e) DIV 2 - no laminin (f) DIV 2 - laminin (g) DIV 3 - no laminin (h) DIV 3 - laminin (i) DIV 10 - no laminin (j) DIV 10 - laminin

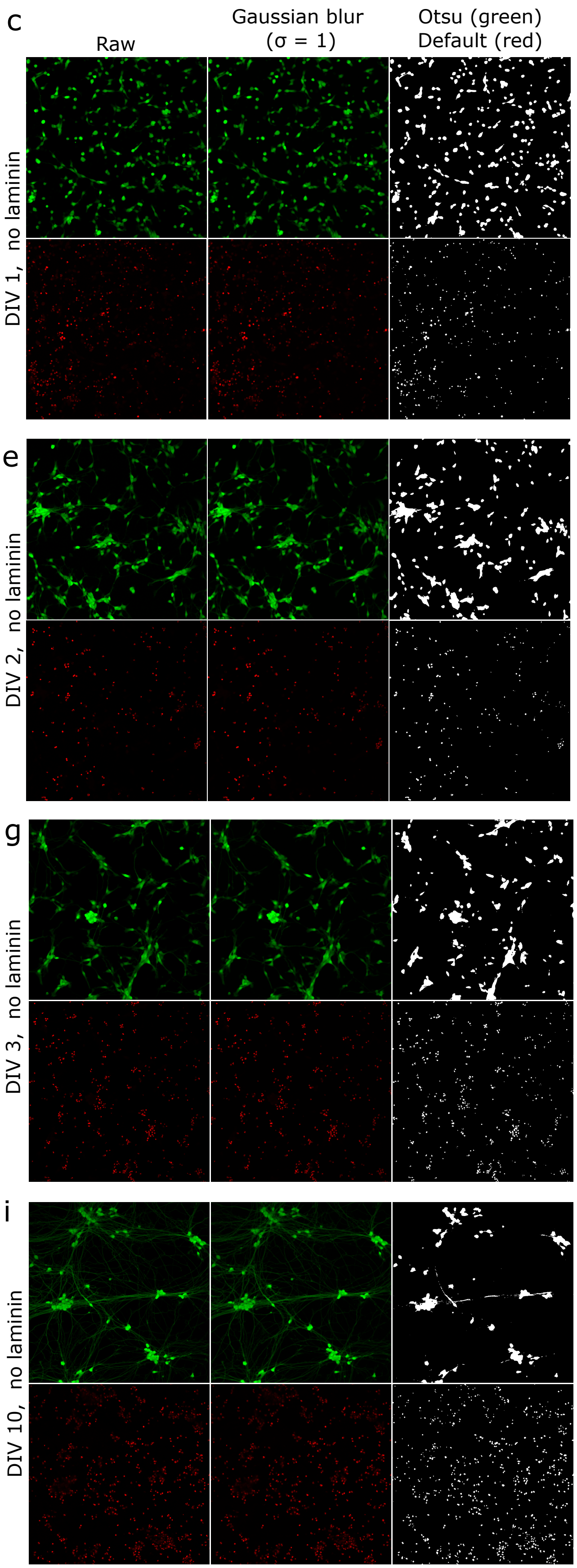


**Figure S 5** See legend on previous page.

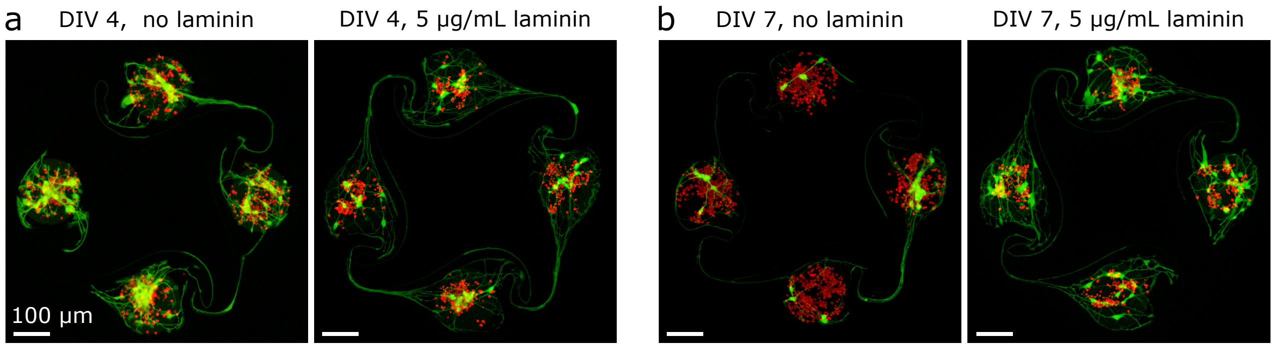


**Figure S 6** Representative examples of fluorescently labelled iNeurons grown in PDMS microstructures at DIV 4 (a) and DIV 7 (b). Green: live cells, stained with CMFDA. Red: dead cells, stained with ethidium homodimer-1.

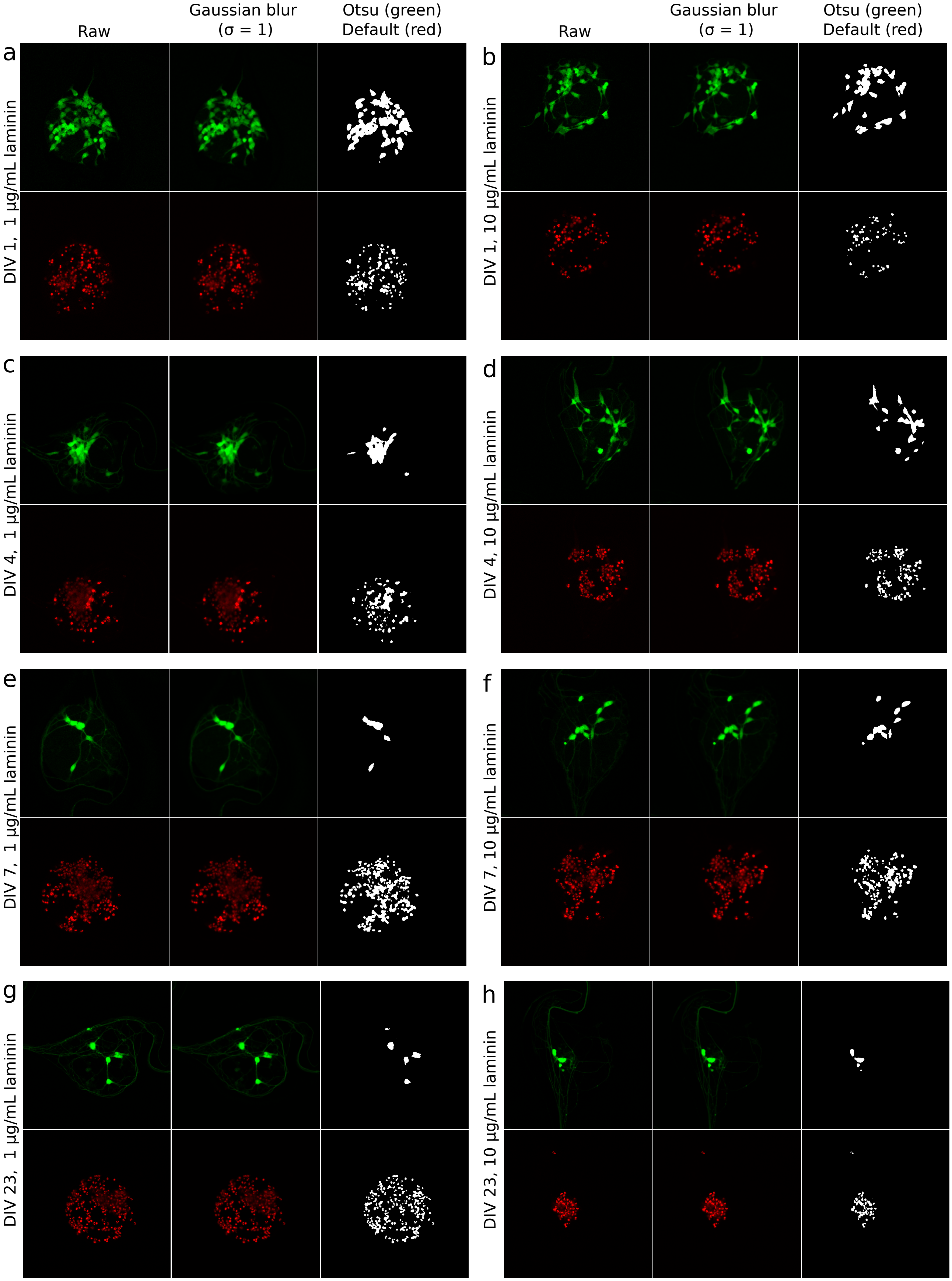


**Figure S 7** Step-by-step image analysis to measure the area occupied by live and dead cells in images of nodes of PDMS microstructures. (a) DIV 1 - 1 $\mu$g/mL laminin (b) DIV 1 - 10 $\mu$g/mL laminin (c) DIV 4 - 1 $\mu$g/mL laminin (d) DIV 4 - 10 $\mu$g/mL laminin (e) DIV 7 - 1 $\mu$g/mL laminin (f) DIV 7 - 10 $\mu$g/mL laminin (g) DIV 23 - 1 $\mu$g/mL laminin (h) DIV 23 - 10 $\mu$g/mL laminin

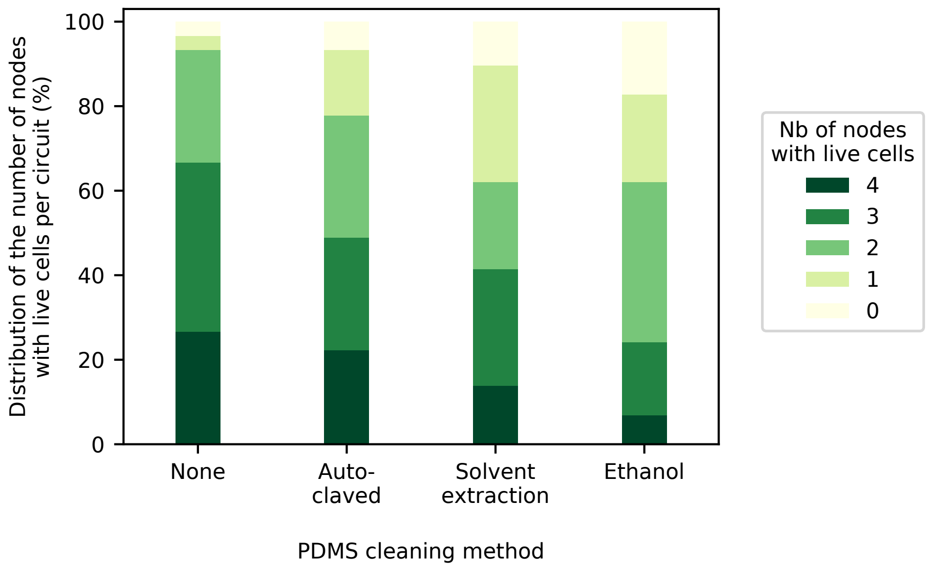


**Figure S 8** Testing the effect of PDMS treatment on the survival of iNeurons in circuits.

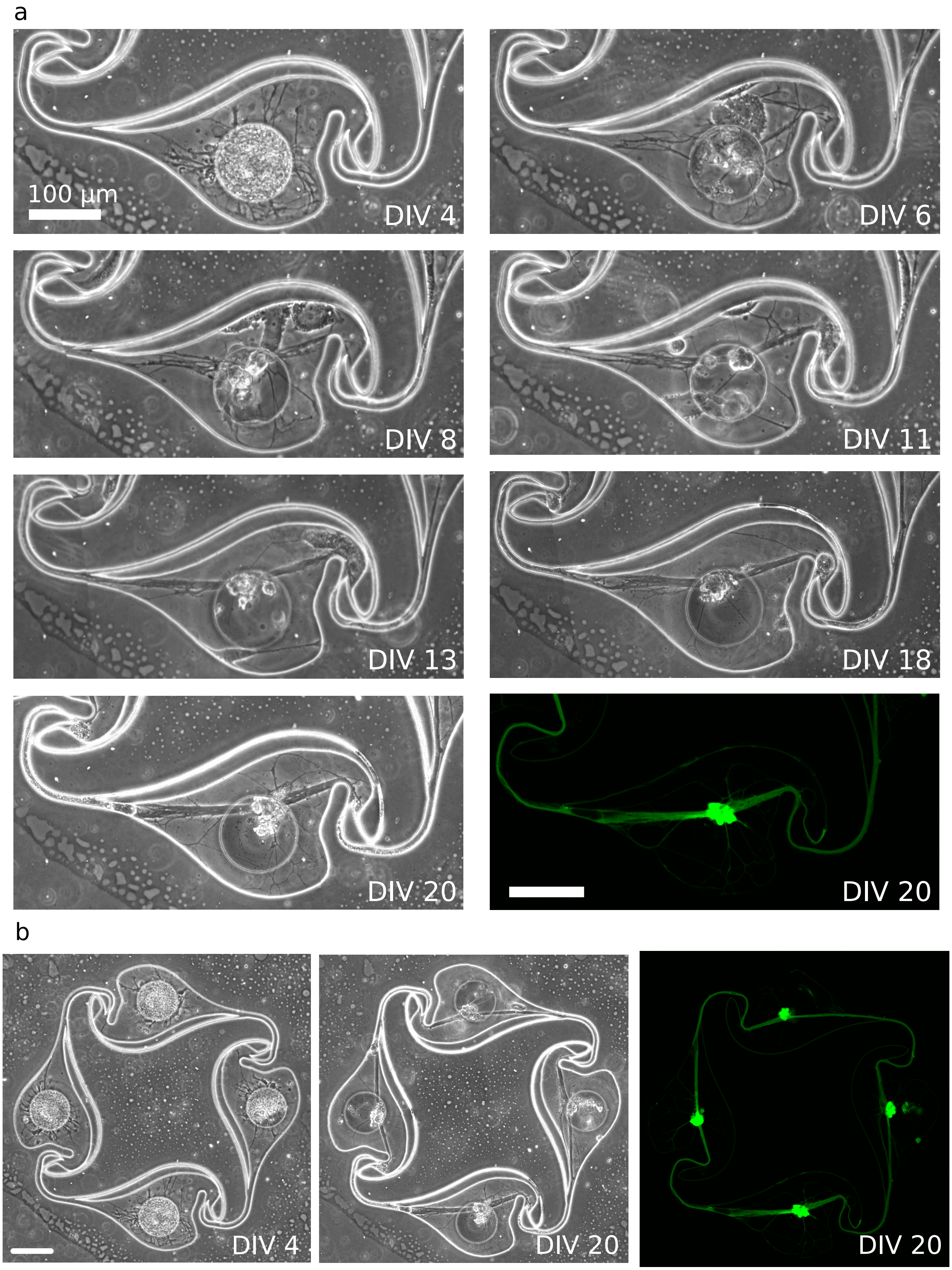


**Figure S 9** Adding macrophages in nodes of iNeurons clears the dead cells. (a) Day-by-day evolution of a node initially filled with many dead iNeurons and where macrophages were added at DIV 4. Images are phase contrast, except for the bottom right one, which shows a Calcein AM fluorescent stain. (b) Full circuit shown in (a) for DIV 4 and DIV 20, with phase contrast and fluorescent Calcein AM stain. Scale bar: 100 $\mu$m.

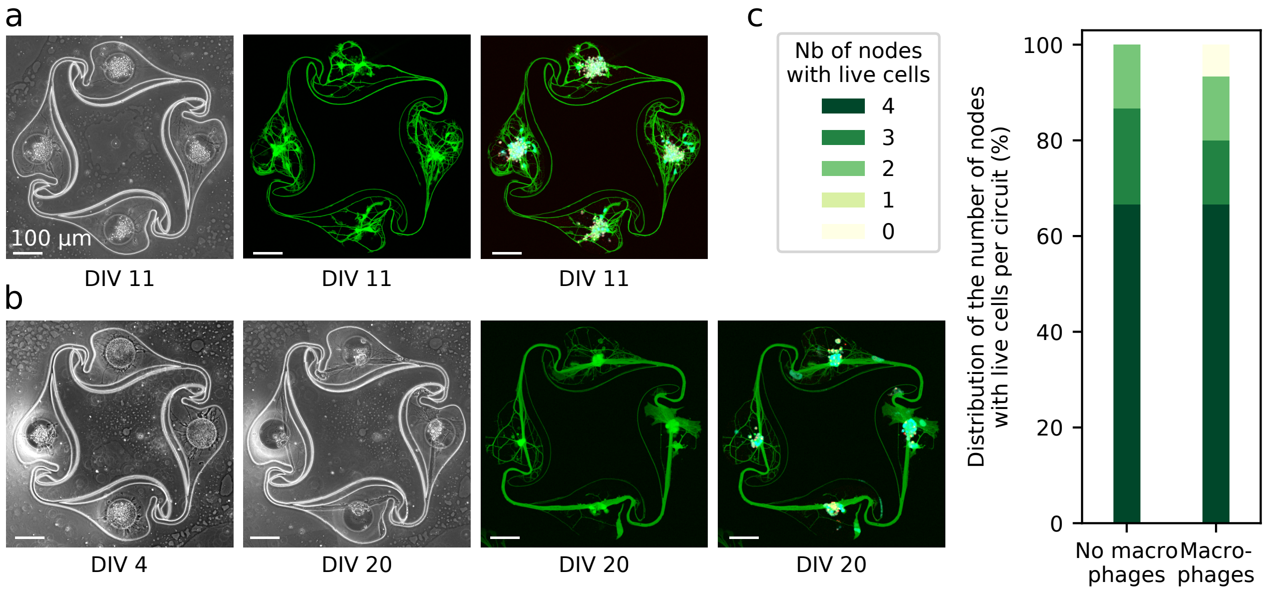


**Figure S 10** Effect of the addition of macrophages on the number of full circuits. (a) Example circuit with iNeurons only, with no macrophages added to the circuit, at DIV 11. (b) Example circuit where macrophages were added to the circuit at DIV 4. Within a few days, the dead cells are cleared by the macrophages. For both (a) and (b): phase contrast pictures (left), Calcein AM stain (center) and overlay of Hoechst, Calcein AM and ethidium homodimer-1 stains (right). (c) Comparing the distribution of nodes with at least one live iNeuron with and without addition of macrophages at DIV 4 (N = 15).

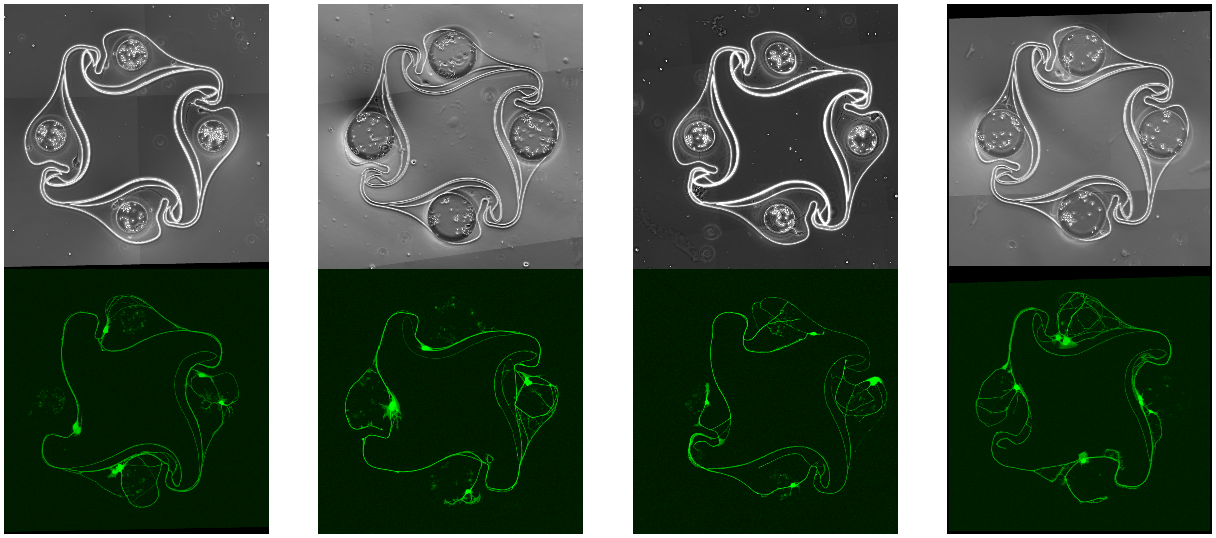

**Condition 1**
No laminin
30 k cells/cm$^2$
100 µm node

**Condition 2**
No laminin
30 k cells/cm$^2$
170 µm node

**Condition 3**
1 µg/mL of laminin
30 k cells/cm$^2$
100 µm node

**Condition 4**
1 µg/mL of laminin
30 k cells/cm$^2$
170 µm node

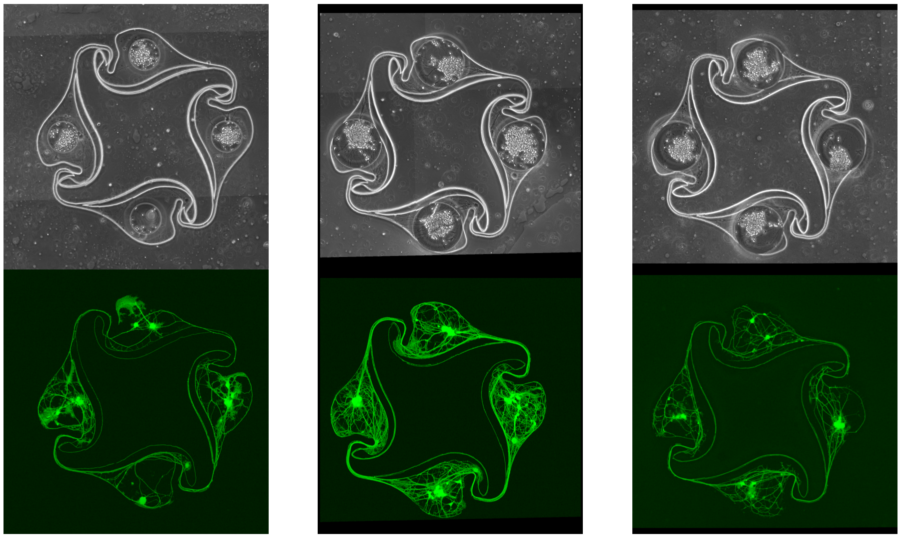

**Condition 5**
1 µg/mL of laminin
65 k cells/cm$^2$
100 µm node

**Condition 6**
1 µg/mL of laminin
65 k cells/cm$^2$
170 µm node

**Condition 7**
10 µg/mL of laminin
65 k cells/cm$^2$
170 µm node

**Figure S 11** Representative examples of circuits with four nodes containing at least one live iNeuron each for the 7 conditions presented in Fig. 5b.

a

b

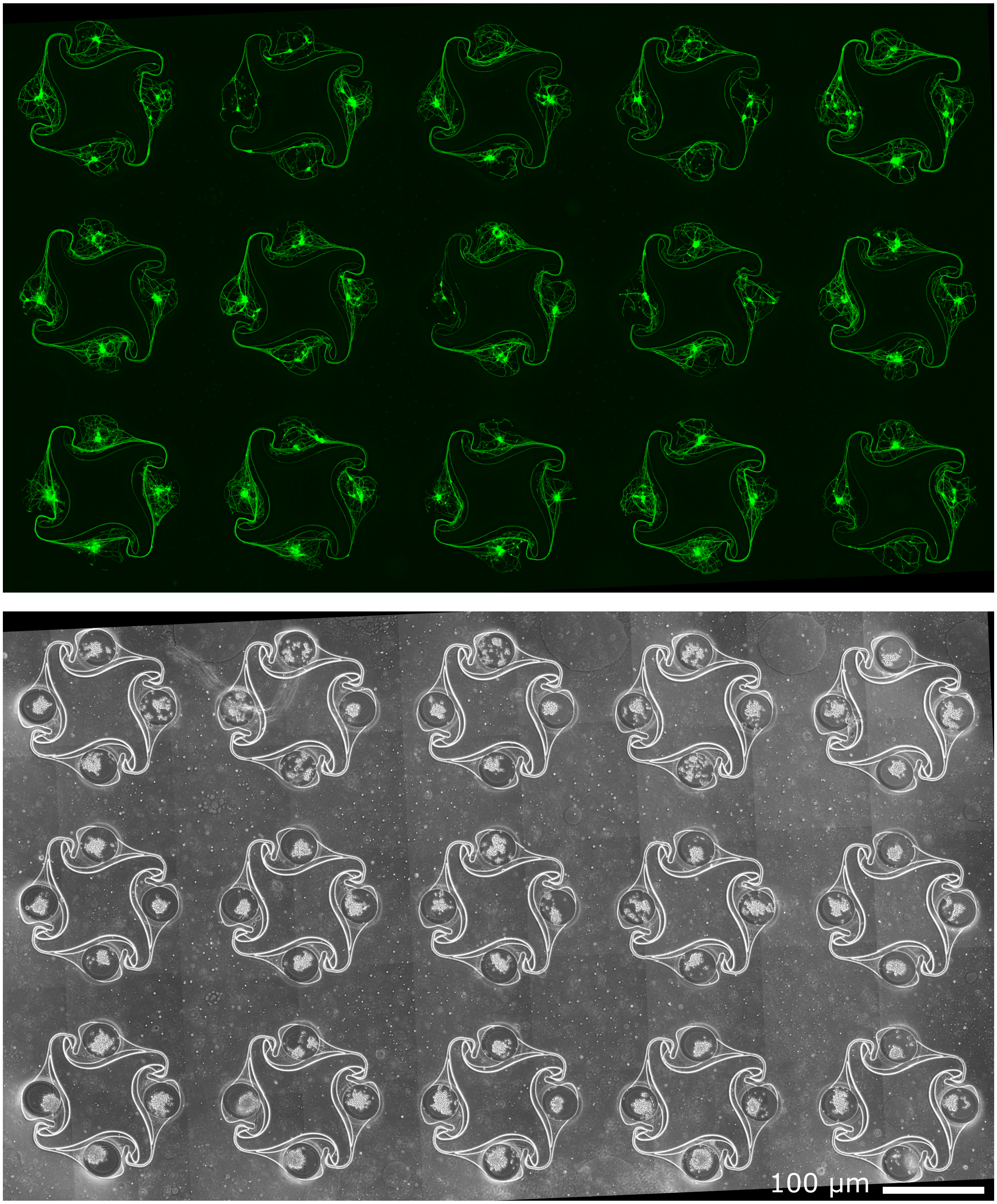


**Figure S 12** Stitched image of the 15 circuits of a PDMS membrane. Top: Calcein AM; bottom: phase contrast. Circuits were seeded at a density of 65k cells/cm$^2$ and 10 $\mu$g/mL of laminin was added to the culture medium during the first week of culture (condition 7 in Fig. S11).

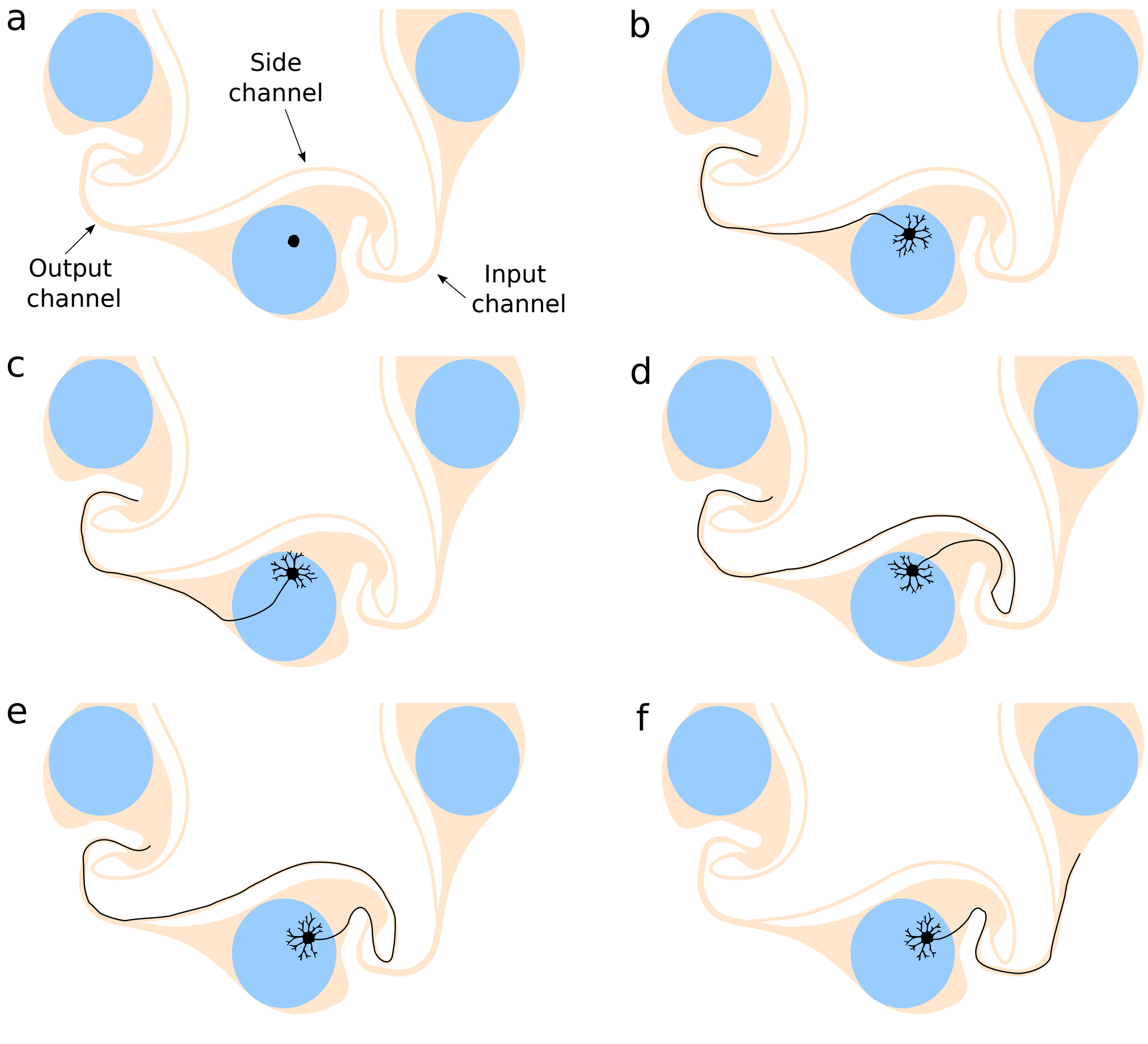


**Figure S 13** Schematics of all the possible directions an axon can grow into after cell seeding. (a) Name of the channels used to describe possible axon outgrowth in the text (with regards to the node where the cell is seeded). Cases (b) to (e) show desirable, clockwise axonal growth direction. Case (f) is the undesirable, counter-clockwise axonal growth direction

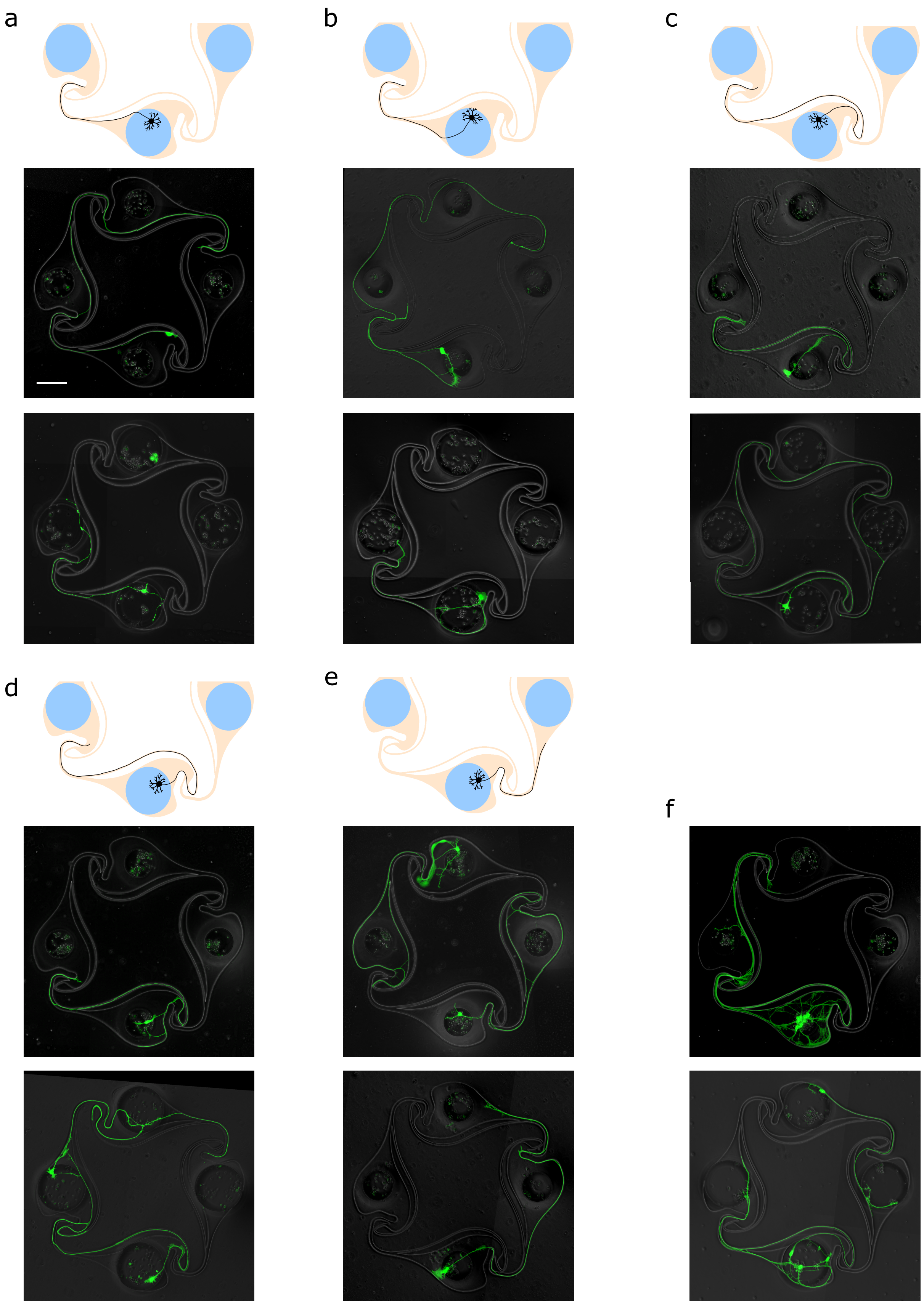


**Figure S 14** Images of circuit with single nodes occupied by iNeurons, illustrating the five possible growth directions for axons in both 100 and 170 $\mu$m diameter nodes (a-e). (f) shows cases where several iNeurons survived in a well and their axons still followed the intended clockwise growth direction. Scale bar: 100 $\mu$m.

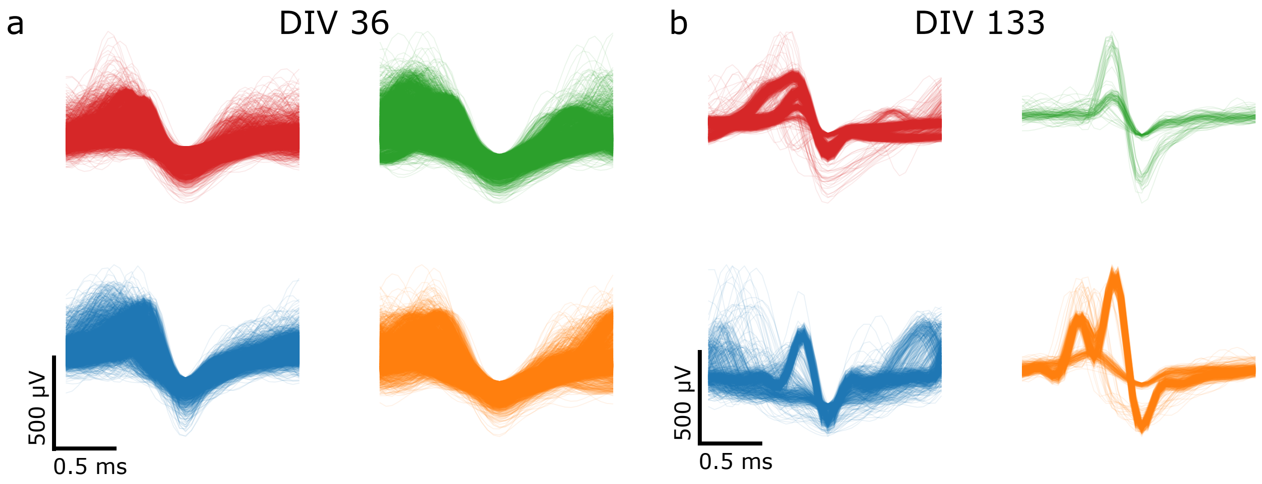


**Figure S 15** Overlay of the waveforms of the action potentials detected in a 5 min recording of spontaneous electrical activity for the four electrodes shown on Fig. 7 at DIV 36 (a) and 133 (b)

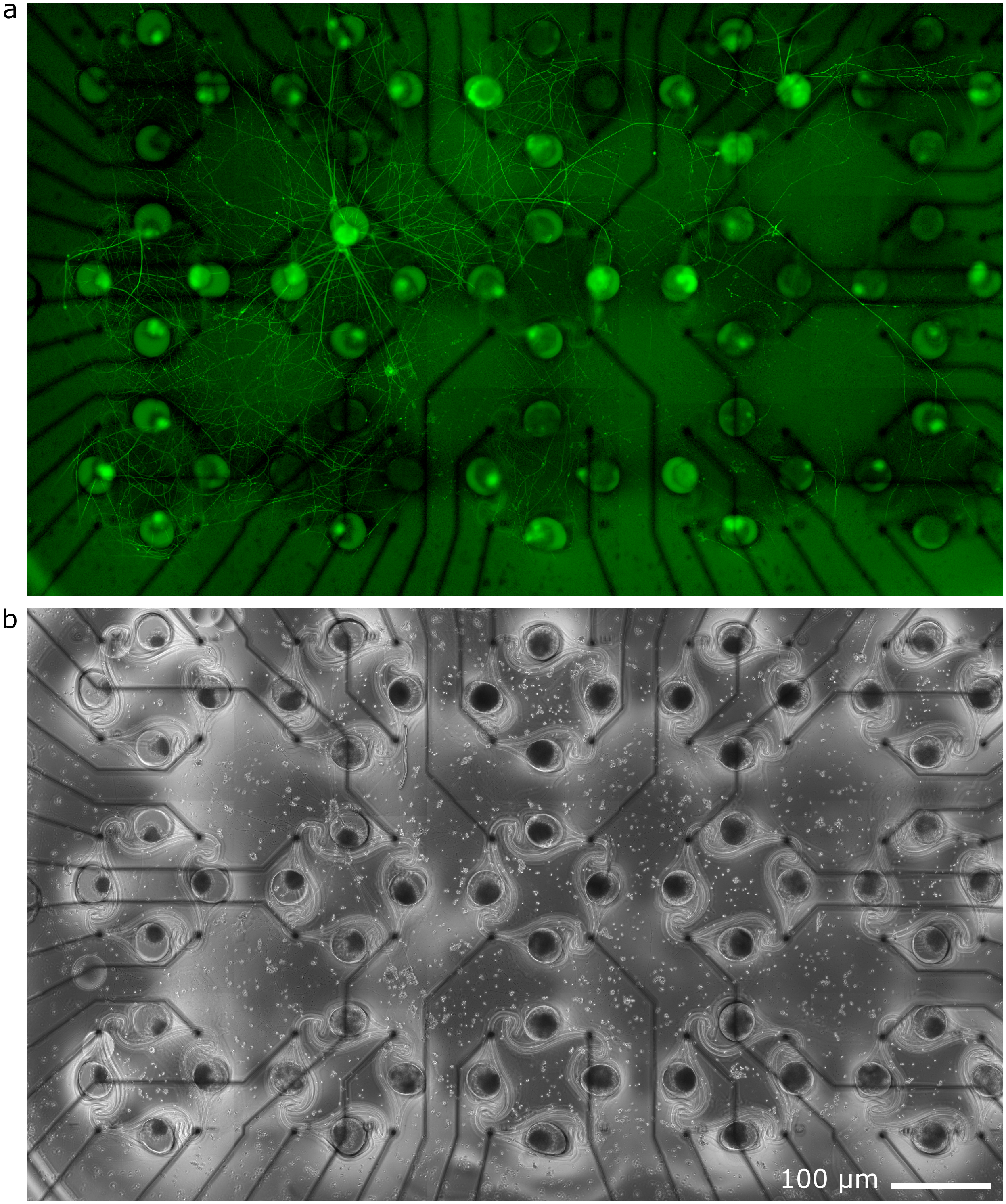


**Figure S 16** Stitched image showing axons growing on top of a PDMS microstructure on a MEA after 91 DIV. (a) CMFDA stain. (b) Phase-contrast image.
